# De novo design of flexible protein interactions with GuideFlip

**DOI:** 10.64898/2026.09.27.754145

**Authors:** Kai Yi, Qingchao Chen, Dongqi Zhang, Pengfei Tian, Jane L. Wagstaff, Stephen H. McLaughlin, Christopher G. Tate, Kiarash Jamali, Sjors H. W. Scheres

## Abstract

De novo design of protein binders requires a target structure. However, for flexible targets, such as intrinsically disordered proteins, this structure does not exist until the binder has stabilized the interaction. Such targets are therefore difficult for methods that separate structure generation from sequence design. We introduce GuideFlip, which co-designs structure and sequence through guided discrete flow matching: binder residues are assigned progressively while the complex is re-predicted at each step, allowing the evolving interface to affect the design process. GuideFlip reduces the hydrophobic bias of direct AlphaFold optimization and improves *in silico* success rates over existing approaches. We release a database of binder candidates for 177 human disordered proteins. Experimentally, we obtain de novo binders to the carboxy terminus of *α*-synuclein and the disordered amino terminus of RBX1 with hit rates of 13.5% and 41.7%, respectively, and we confirm the epitopes of selected binders by NMR and mutagenesis. Applying GuideFlip to flexibility on the binder side, we design a nanobody that binds the agonist-bound *β*_1_-adrenergic receptor in its active state, but not the receptor in its inactive state, with a 75% hit rate and a cryo-EM structure confirming the design. GuideFlip enables protein design where bound structures emerge only upon binding.

## Introduction

Conformational flexibility is a common strategy for achieving specific molecular recognition in biology^1^. It broadens the range of geometries accessible to a binding surface and enables induced fit and conformational selection, while allowing sequence diversity to be concentrated in regions that can evolve without compromising the stability of a surrounding scaffold^2,3^. Flexible interactions have important roles in cellular signalling^4^, immunity, and infection^5^, and many are of therapeutic relevance^6^. Antibodies and T-cell receptors use flexible loops with diverse sequences to recognize structurally heterogeneous targets^7^, whereas many regulatory proteins recognize short linear motifs in intrinsically disordered regions that only acquire structure upon binding^8^. The ability to engineer such interactions would extend the range of medically relevant molecular recognition problems accessible to protein design^9^.

However, most methods for computational binder design work best when the interaction is represented by a well-defined structure. For such cases, advances in machine learning have enabled the routine creation of high-affinity binders^10^ using a sequential strategy: first generating a backbone that forms a geometrically-complementary interface with a target^11,12^ and then designing a sequence compatible with that backbone^13^. But the separate design of backbone structure and sequence has proven more difficult for interactions that involve conformational flexibility, because mutations in the binder alter its conformational ensemble at the same time as they alter the energetics of the desired interface. Recently, methods aimed at the design of binders to intrinsically disordered regions have been proposed^14,15^, but their success depends on prior structural knowledge.

Backpropagation through differentiable structure prediction models provides one way of coupling sequence and structure during design^16,17^. BindCraft^18^, for example, optimizes binder sequences through AlphaFold2 (AF2)^19^, allowing the structures of the target and binder to fold together throughout optimization. However, AlphaFold was trained to predict protein structures, not to generate protein sequences. Direct optimization of its confidence metrics can therefore produce sequences that differ markedly from those of natural proteins. One manifestation of this is a bias towards hydrophobic residues^20^, which favour dense packing and high-confidence structures. BindCraft addresses this problem by using SolubleMPNN^13^ to redesign sequences after backpropagation. By this stage, however, the backbone and much of its packing environment have already been selected. Instead, combining structural gradients with a generative model of discrete protein sequence would allow sequence and structure to be optimized together throughout the design process.

Here we introduce GuideFlip, a framework for protein design that jointly optimizes sequence and structure through guided discrete flow matching (Fig. 1). We combine all-atom inverse folding^21^ with structural gradients obtained by backpropagation through AlphaFold-Multimer (AF-Multimer)^22^. Rather than separating structure optimization from sequence design, AF-Multimer gradients guide the discrete generative flow through sequence space, coupling the search for plausible protein sequences to the conformations and interactions they encode. This allows sequence and structure to adapt together throughout design. We apply this framework to challenging recognition problems involving conformational flexibility, obtaining *de novo* binders against the carboxy-terminal region of *α*-synuclein and the disordered amino-terminal region of RBX1 with experimental hit rates of 13.5% and 41.7%, respectively. On the binder side, we design the complementarity-determining regions (CDRs) of nanobodies targeting the active conformation of the *β*_1_-adrenergic receptor, achieving an experimental hit rate of 75%. These results establish guided discrete flow matching as a general framework for designing protein interactions through the concurrent optimization of sequence and structure.

**Figure 1:**
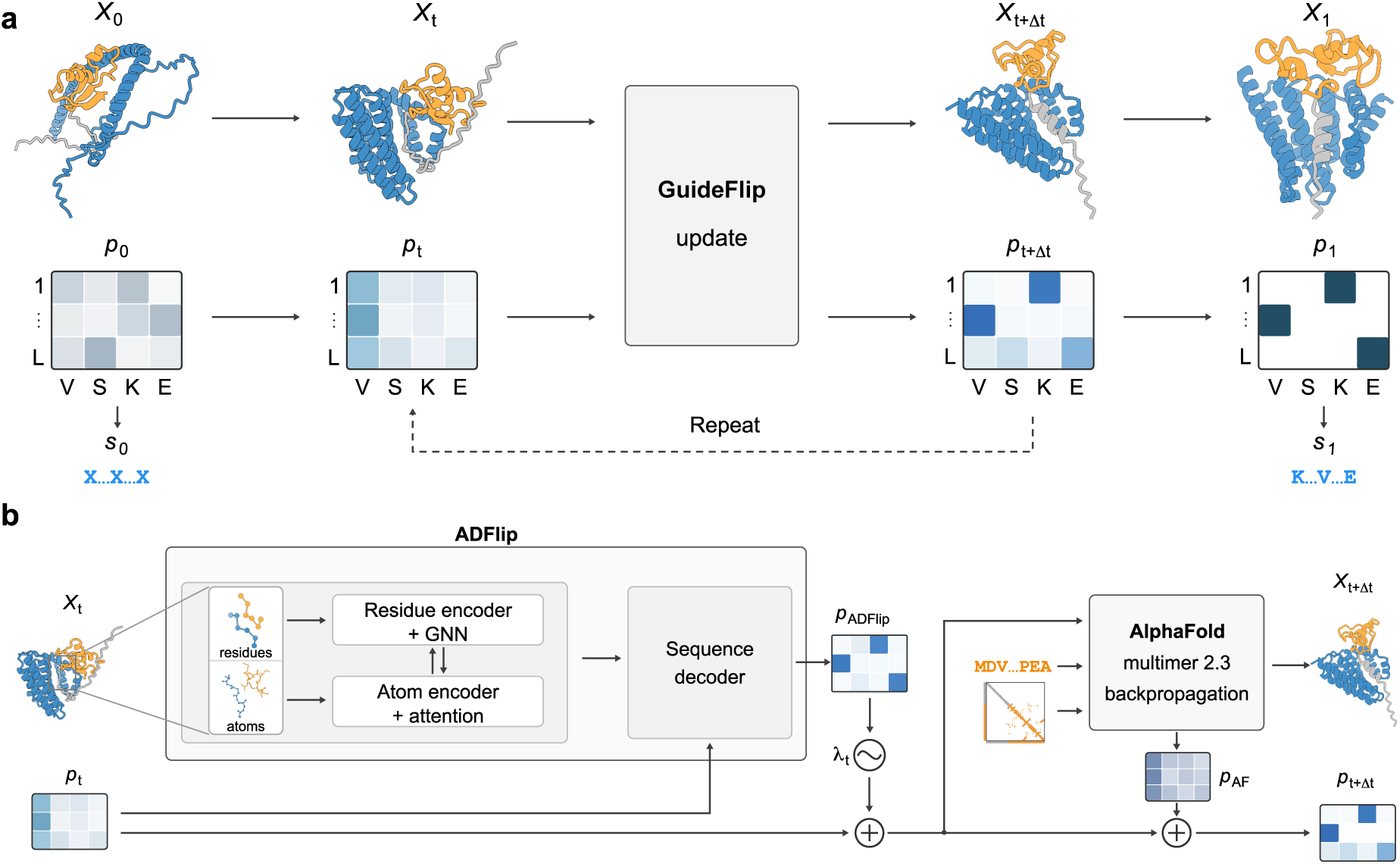
The GuideFlip design framework. **a**, GuideFlip jointly updates the binder sequence distribution *p_t_* and predicted target–binder structure *X_t_*through guided discrete flow matching (detailed in **b**). Generation proceeds from random initialization (*p*_0_) to a one-hot distribution (*p*_1_) specifying the final binder sequence. Matrices schematically represent amino-acid distributions: columns denote binder positions (*L*, binder length), and rows show example amino-acid identities. **b,** At each step, GuideFlip takes *X_t_* and *p_t_*as inputs. ADFlip combines residue- and atom-level structural encodings with sequence information from *p_t_* to predict an amino-acid prior, *p*_ADFlip_. The contribution of *p*_ADFlip_ is controlled by *λ_t_*and omitted when the structure is insufficiently confident. Combining this prior with *p_t_* supplies the binder input to AlphaFold-Multimer, alongside the target sequence and structural template (orange). Templates are retained for rigid regions and omitted for flexible regions. AlphaFold predicts the updated complex *X_t_*_+Δ*t*_. Backpropagation of the design loss yields gradients with respect to the binder input probabilities, which are used to approximate the guidance distribution *p*_AF_. Combining *p_t_*, *p*_ADFlip_ and *p*_AF_ gives *p_t_*_+Δ*t*_. Structures are predicted models, with binder in blue and target in orange and grey.

## Designing flexible interactions with GuideFlip

GuideFlip uses guided discrete flow matching to jointly design binder sequences and target–binder complexes (Fig. 1). Starting from random amino-acid probabilities, GuideFlip progressively assigns amino-acid identities to initially masked binder positions, while updating the predicted complex. The binder sequence and complex conformation are therefore explored together during generation.

GuideFlip combines two sources of information to guide this process. ADFlip^21^, our all-atom inverse folding model, predicts amino-acid preferences from the current complex and partially specified binder sequence. Its residue- and atom-level encoders combine information about the backbone geometry and local atomic environment, including target side chains. To support the design of soluble binders, we retrained ADFlip on experimentally determined structures from the Protein Data Bank, excluding membrane proteins, following a strategy similar to that used for SolubleMPNN^13^. The model supplies a structure-conditioned sequence prior.

Backpropagation through AF-Multimer^22^ provides structural feedback during sequence generation. We combine the ADFlip prior with the current amino-acid probabilities and supply them to AF-Multimer. AF-Multimer predicts the complex and provides confidence and structural measures for a design loss. Backpropagation of this loss adjusts the amino-acid preferences, and the updated probabilities and predicted complex are carried into the next round.

Like BindCraft, GuideFlip uses backpropagation through AF-Multimer to optimize binder sequences. BindCraft obtains discrete sequences through a staged optimization procedure that includes temperature annealing, followed by SolubleMPNN redesign of non-interface residues. GuideFlip instead introduces a structure-conditioned sequence prior from ADFlip once an initial binder structure is obtained. This prior is combined with AF-Multimer gradients to progressively resolve amino acids, without temperature annealing or a separate SolubleMPNN redesign stage. Learned sequence preferences therefore guide interface residue selection while the sequence and predicted complex are still evolving.

Structural templates control which regions retain input structural information and which can explore alternative conformations. For disordered targets, templates are omitted from flexible regions, allowing their structure to change with the emerging binder. For antibodies, framework templates are retained, whereas the CDRs are redesigned without template constraints. This allows the same framework to address both disordered target regions and flexible binding loops, such as those in nanobodies.

### Designing binders for human disordered proteins

We compared GuideFlip with RFdiffusion^11^ and BindCraft^18^ on ten targets with experimentally validated binders (Fig. 2). These comprised eight targets from a published study of binder design against intrinsically disordered regions^14^, together with *α*-synuclein and RBX1, for which we experimentally validated binders in this study. For each method, we evaluated 1,000 designs per target using the same combined AlphaFold3 (AF3)^23^ filters used to select candidates for experimental testing. GuideFlip yielded a higher computational success rate than both alternative methods on eight of the ten targets (Fig. 2a,b).

**Figure 2:**
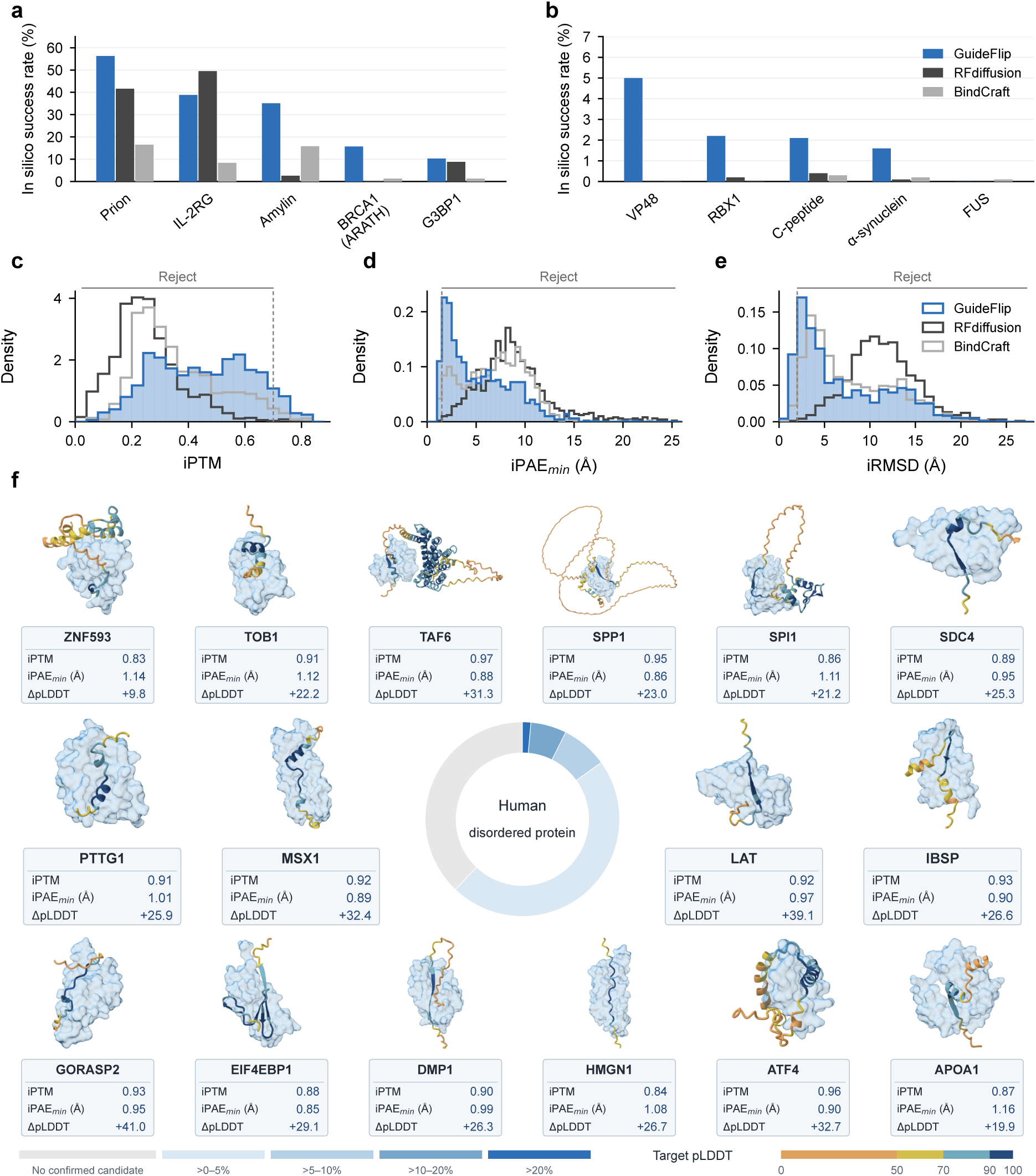
*In silico* benchmarking and target coverage of GuideFlip. **a,b**, Computational success rates for ten disordered protein targets, plotted on different y-axis scales. Blue, dark grey and light grey denote GuideFlip, RFdiffusion and BindCraft, respectively. Designs pass if they meet all three criteria: iPTM *≥* 0.7, iPAE_min_ *≤* 1.5 Å and iRMSD *≤* 2 Å. Each method was run to produce 1,000 designs per target. Success rates use all 1,000 designs as the denominator, including designs without successful evaluation, which count as non-passing. **c**–**e**, Distributions of AlphaFold3 interface predicted TM score (iPTM; **c**), minimum interface predicted aligned error (iPAE_min_; **d**) and interface root-mean-square deviation (iRMSD; **e**) for binder designs targeting the carboxy terminus of *α*-synuclein. Vertical dashed lines (grey) mark filtering thresholds for successful designs. **f**, Target coverage for 285 human protein targets selected using annotated disorder content greater than 30% in DisProt and protein length below 700 amino acids. At least one design met all three criteria for 177 of the 285 targets (62.1%). Representative AlphaFold3-predicted complexes are shown. Binders are shown as light-blue surfaces and targets as ribbons coloured by predicted local distance difference test (pLDDT) scores. Each card reports iPTM, iPAE_min_ (Å) and target-interface ΔpLDDT, calculated as complex minus apo pLDDT over the same target interface residues. Target-interface residues have at least one heavy atom within 5 Å of a binder heavy atom in the predicted complex.

We examined each filtering metric separately for the *α*-synuclein designs. Compared with both alternative methods, the distributions for GuideFlip designs were shifted towards higher AF3 interface predicted TM score (iPTM) and lower minimum interface predicted aligned error (iPAE_min_; Fig. 2c,d). GuideFlip designs also showed lower interface root-mean-square deviation (iRMSD) between the design models and AF3 predictions, indicating closer agreement at the binding interface (Fig. 2e).

To assess the scalability of our approach, we curated all human intrinsically disordered proteins (*n* = 285) catalogued in the DisProt database^24^. We chose proteins with fewer than 700 residues, of which more than 30% were annotated as disordered. We used per-residue predicted local distance difference test (pLDDT) scores from AF3 predictions to select short, putatively disordered fragments, and generated 500 binder designs per target. Designed binder sequences were co-folded with their full-length targets using AF3 and filtered by iPTM, minimum interface PAE and interface RMSD. Overall, at least one computational candidate passed all three filters for 177 of the 285 targets (62.1%; Fig. 2f). We distribute the successful designs as a database^1^.

To validate these *in silico* results, we chose two human disordered proteins to experimentally assess our designed binders: *α*-synuclein and RBX1.

### De novo binders against the carboxy terminus of *α*-synuclein

*α*-synuclein is an intrinsically disordered protein with an amino-terminal region, a hydrophobic central region and a negatively charged C-terminal region. Its aggregation into amyloid filaments is a pathological hallmark of Parkinson’s disease, dementia with Lewy bodies, multiple system atrophy and other synucleinopathies^25–28^. The C-terminal region remains exposed upon filament formation^29–31^ and has a potential role in regulating *α*-synuclein aggregation, making it an attractive target for diagnostic and therapeutic binders. However, this region is challenging for structure-based protein design because it has never been observed in a folded conformation. Conventional structure-based design methods therefore have no fixed backbone against which to design.

We used GuideFlip to design binders against *α*-synuclein residues 100–140 (Fig. 3a) without a structural template. We specified no binding hotspots, allowing AF-Multimer to select the residues involved in binding. Following *in silico* filtering, we experimentally tested binding for 52 sequences (Supplementary Table S1). Of these, 37 showed good expression levels and were further tested by surface plasmon resonance (SPR), using biotinylated full-length *α*-synuclein immobilized on a streptavidin sensor chip. Because almost all lysines in *α*-synuclein lie within its amino-terminal and hydrophobic regions, this immobilization leaves the C-terminal region accessible for binding. Seven of the 37 designs exhibited detectable binding (Extended Data Fig. 1; Supplementary Table S2). The protein designs dnAS15 (Fig. 3b,c) and dnAS36 showed the highest affinities, with *K*_D_ values of 211 and 566 nM, respectively (Extended Data Fig. 1a,b).

**Figure 3:**
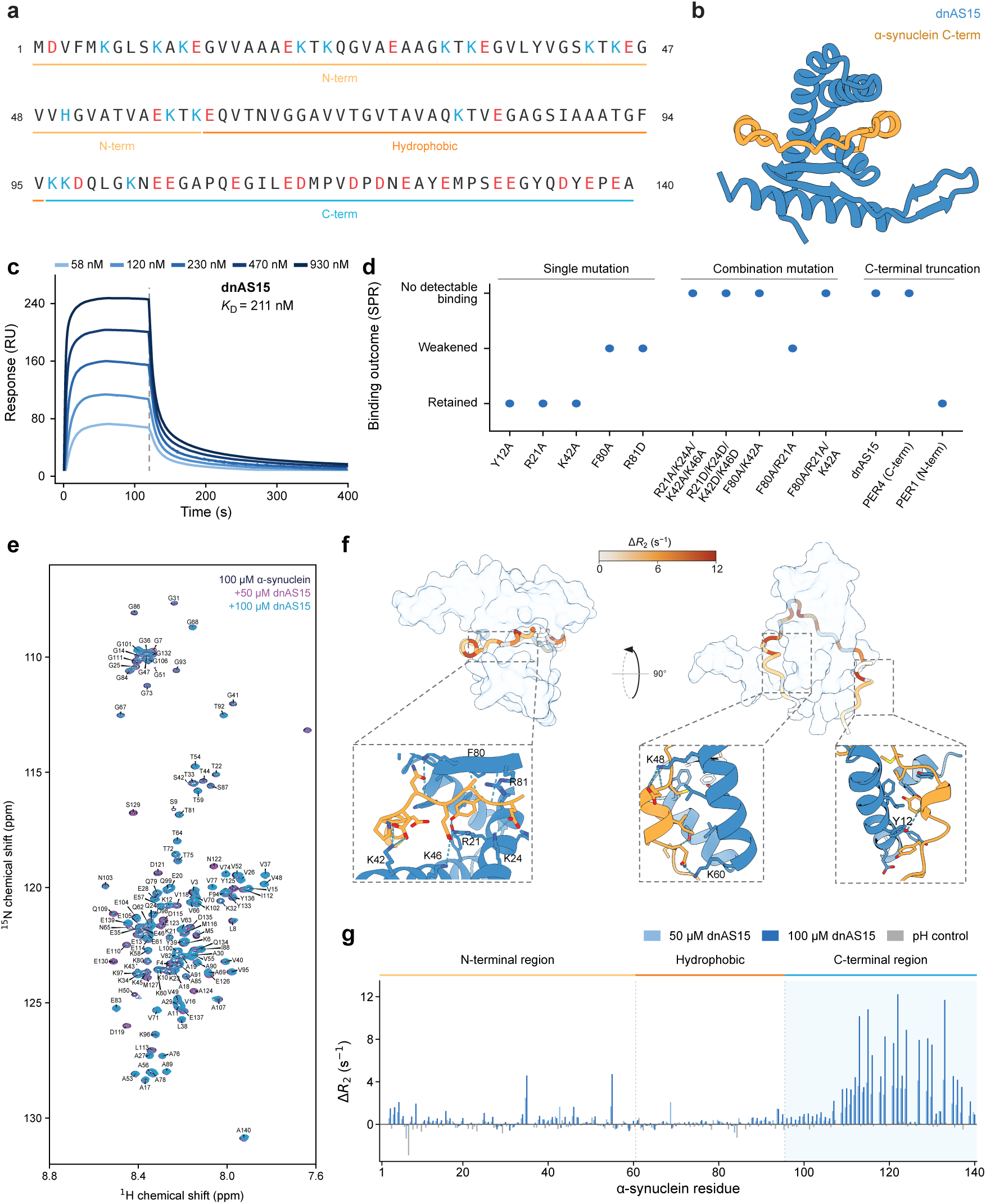
*De novo* binders against the disordered C terminus of *α*-synuclein. **a**, *α*-synuclein sequence, with basic residues in blue, acidic residues in red and the amino-terminal (N-term), hydrophobic and C-terminal (C-term) regions indicated. **b**, Predicted dnAS15–*α*-synuclein complex. dnAS15 is shown in blue and *α*-synuclein residues 105–140 in orange. **c**, SPR sensorgrams of dnAS15 binding to full-length *α*-synuclein (*K*_D_ = 211 nM). **d**, Binding effects of dnAS15 mutations and *α*-synuclein truncation, assessed by SPR. The first two groups show dnAS15 variants binding to full-length *α*-synuclein; the third shows binding to the C-terminal truncated construct (residues 1–121, lacking the final 19 residues). PER4 and PER1 recognize C-terminal and amino-terminal epitopes, respectively. **e**, Overlay of two-dimensional ^1^H–^15^N HSQC spectra of 100 *µ*M full-length *α*-synuclein alone and with 50 or 100 *µ*M dnAS15. Assigned backbone amide peaks are labelled. **f**, Orthogonal views of the predicted complex. dnAS15 is shown as a surface; *α*-synuclein is coloured by NMR-measured changes in backbone transverse relaxation rates (Δ*R*_2_) for 100 *µ*M *α*-synuclein with 100 *µ*M dnAS15, relative to *α*-synuclein alone. Unassigned residues are grey. Boxed regions are enlarged below, with dnAS15 in blue and *α*-synuclein in orange. Selected dnAS15 residues are labelled; cyan dashed lines indicate putative hydrogen bonds and salt bridges. **g**, Per-residue Δ*R*_2_ for 100 *µ*M full-length *α*-synuclein with 50 or 100 *µ*M dnAS15, relative to *α*-synuclein alone. Grey bars show the pH control (*α*-synuclein with 1 *µ*l of 1 M HCl). Bars show point estimates for 131 assigned backbone amides; the C-terminal region is shaded.

To test whether dnAS15 recognizes the intended C-terminal epitope, we compared its binding to full-length *α*-synuclein and a C-terminal truncated construct lacking the final 19 residues (122–140) by SPR. dnAS15 bound full-length *α*-synuclein but showed no detectable binding to the truncated construct; the control antibody PER1^29^, which recognizes residues 11–34, bound both constructs, whereas PER4^29^, which recognizes the C-terminal region, bound only full-length *α*-synuclein (Fig. 3d).

In the predicted complex with dnAS15, *α*-synuclein residues 107–139 wrap around the binder in a largely extended conformation interrupted by two short *α*-helices (Fig. 3b). To assess the designed binding mode, we mutated dnAS15 residues predicted to contact *α*-synuclein and measured binding by SPR (Fig. 3d,f; Supplementary Figs. S1, S2). Single-residue mutations retained or weakened binding, whereas four of five tested combinations of mutations at distinct interfaces abolished detectable binding.

To confirm the designed binding sites, we also performed solution-state nuclear magnetic resonance (NMR) measurements using ^15^N-labelled full-length *α*-synuclein and unlabelled dnAS15 (Fig. 3e–g; Extended Data Fig. 2) or dnAS36 (Extended Data Fig. 3). For both binders, backbone transverse relaxation rates (*R*_2_) increased markedly in the C-terminal region, whereas the amino-terminal and hydrophobic regions showed little change. Increased transverse relaxation rates upon binder addition are consistent with involvement of the affected *α*-synuclein residues in binding.

These results suggest that both binders recognize the C-terminal domain of *α*-synuclein as designed.

### De novo binders against the disordered tail of RBX1

RBX1, which is also known as ROC1, is the RING component of several cullin–RING ubiquitin ligases^32^. Its N-terminal region is disordered in isolation^33^, but folds into an intermolecular *β* -sheet upon binding to the cullin scaffold, forming part of the catalytic ligase core^34^. RBX1 thus provides a biologically relevant example of molecular recognition coupled to a disordered-to-ordered transition. Cullin-RING ligase activity controls the turnover of numerous regulators of cell proliferation and signaling^32^ and has been extensively pursued therapeutically, particularly in cancer^35,36^.

We tested whether GuideFlip could design binders against the disordered N-terminal region of RBX1 (Fig. 4a). During design, we omitted structural template information for RBX1 residues 1–40, allowing this region to change conformation, and designated residues 28–38 as binding hotspots. We generated 100 sequences that passed our *in silico* filter and submitted seven for experimental testing by Adaptyv Bio in the GEM *×* Adaptyv RBX1 binder-design competition^37^. Surface plasmon resonance (SPR) identified two binders, dnRB1 and dnRB2, with affinities (*K*_D_) of 85 nM and 170 nM, respectively (Fig. 4b,c; Extended Data Fig. 4). dnRB1 showed slower association and dissociation than dnRB2 (Extended Data Fig. 4). In a second design round, we tested five additional designs by biolayer interferometry (BLI), three of which bound RBX1. The strongest binder, dnRB8, had a *K*_D_ of 4.5 nM (Extended Data Fig. 4; Supplementary Tables S3, S4).

**Figure 4:**
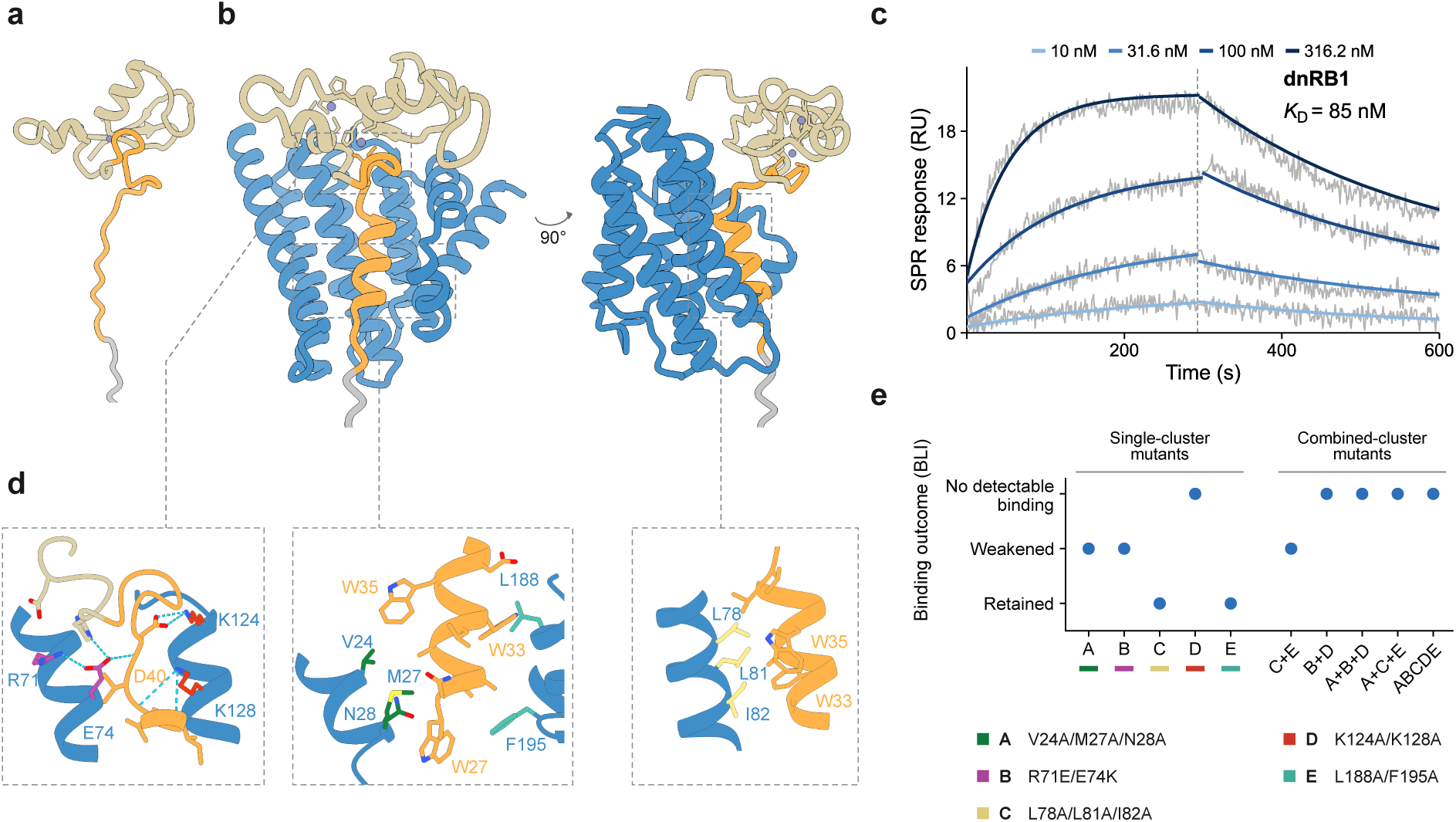
A *de novo* binder to the disordered N-terminal region of RBX1. **a**, Model of RBX1. **b,** Orthogonal views of its predicted complex with dnRB1. RBX1 colours indicate the targeted N-terminal segment (orange), remaining N-terminal residues (grey) and RING domain (beige). dnRB1 is blue; purple spheres represent zinc ions. **c**, Surface plasmon resonance (SPR) sensorgrams of RBX1 binding to immobilized dnRB1. Grey traces show measured responses; blue curves show global 1:1 binding-model fits. The dashed line marks dissociation onset. **d**, Close-ups of the predicted interface boxed in **b**. Side-chain colours identify mutation clusters in **e**; residue labels identify dnRB1 (blue) and RBX1 (orange). Cyan dashed lines indicate predicted hydrogen bonds. **e**, BLI binding outcomes for dnRB1 interface mutants relative to wild-type dnRB1. Each point represents one construct, classified as retained, weakened or no detectable binding under the assay conditions. Single-cluster substitutions are shown below the plot. B+D and A+B+D contain R71A/E74A, whereas B contains R71E/E74K. ABCDE contains M27A/E74K/L78A/K128D/L188A.

To assess the accuracy of the designed binding mode, we introduced mutations at the dnRB1–RBX1 interface. In the designed complex, RBX1 residues 29–38 form an *α*-helix that is encircled by dnRB1, which also packs against the surface of the RING domain (Fig. 4b,d). We mutated five clusters of interface residues in dnRB1 (A–E) and assessed binding by BLI (Fig. 4d,e). Mutating clusters A or B weakened binding, whereas mutating C or E alone had no detectable effect. Mutating cluster D abolished binding, identifying a dominant hotspot within the distributed interface. Combined mutations of C and E weakened binding, whereas all tested combinations involving cluster D abolished it. Simultaneously mutating clusters A, C and E also abolished binding (Supplementary Figure S3). The close correspondence between mutational sensitivity and the designed contacts suggests that dnRB1 engages RBX1 as designed.

### Designing conformation-specific GPCR nanobodies

In the case of binders against *α*-synuclein and RBX1, the flexibility was limited to the target. With GuideFlip, we can also allow regions of flexibility in the binder, while fixing the target in a chosen conformation. Natural instances of this are the interactions that antibodies and nanobodies make with their targets through their complementarity-determining region (CDR) loops.

We therefore designed nanobody CDRs to recognize a specific conformation of a G protein-coupled receptor (GPCR). GPCRs are major drug targets, but their conformational dynamics complicate structural studies. Intracellular nanobodies can bind the cytoplasmic cleft and selectively stabilize defined receptor conformations, facilitating structure determination^38–41^. This conformational selectivity also enables screening for small molecules that preferentially bind particular receptor states^42^. However, obtaining such nanobodies remains challenging and has often relied on animal immunization. With GuideFlip, we targeted the active state of the *β*_1_-adrenergic receptor (*β*_1_AR), a prototypical cardiovascular drug target (Fig. 5a).

**Figure 5:**
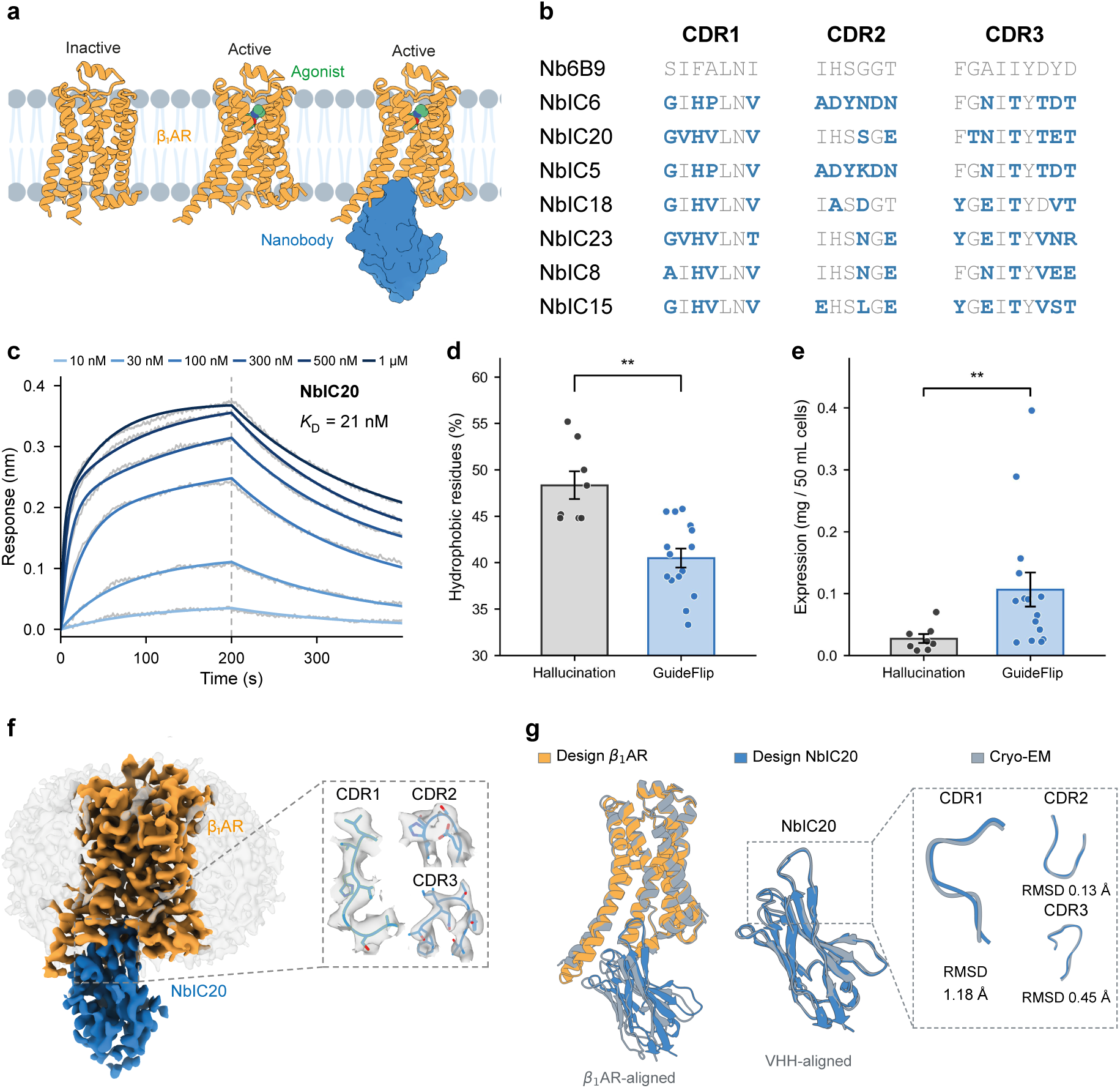
Intracellular nanobodies against active-state *β*_1_AR. **a**, Schematic of *β*_1_AR in its inactive, agonist-bound active and nanobody-bound active states. **b**, Complementarity-determining region (CDR) sequences of Nb6B9 and seven GuideFlip designs. Residues identical to Nb6B9 are grey; substitutions are shown in bold blue. **c**, Biolayer interferometry sensorgrams for NbIC20 binding to isoprenaline-bound *β*_1_AR at six concentrations (10–1,000 nM). Grey traces show measured responses; blue curves show global 1:1 binding-model fits (*K*_D_ = 21 nM), with darker colours indicating higher concentrations. The dashed line marks dissociation onset. **d,e**, Predicted interface hydrophobicity (**d**) and soluble expression yield in *E. coli* (**e**) for GuideFlip (blue; *n* = 15) and hallucination-only designs (grey; *n* = 8). Points represent individual designs; bars and error bars show the mean and standard error of the mean, respectively. *^∗∗^P <* 0.01, two-sided Mann–Whitney U test. **f**, Cryo-EM reconstruction of the isoprenaline-bound *β*_1_AR–NbIC20 complex at 3.3 Å resolution, with *β*_1_AR in orange and NbIC20 in blue. Insets show CDR models and density (grey). **g**, Design model (*β*_1_AR, orange; NbIC20, blue) overlaid with the cryo-EM structure (blue-grey), aligned on *β*_1_AR (left) or VHH (centre). Insets compare CDR conformations, with RMSDs indicated.

We used the X-ray structure of the well-characterized Nb6B9 nanobody bound to the active state of the *β*_1_-adrenergic receptor as a scaffold (PDB: 6H7M)^38,43^ and used GuideFlip to design its three CDRs *de novo* (Fig. 5b). We kept both the variable domain of the heavy chain of the heavy-chain antibody (VHH) framework and the active-state *β*_1_AR fixed during design and targeted receptor hotspot residues Arg91, Ile95, and Ala212 in the intracellular cleft. For each of 200 design trajectories, GuideFlip generated two sequences. We kept trajectories only when all their sequences passed the filter. This yielded 16 designs from eight trajectories, with sequences distinct from that of Nb6B9 (Supplementary Table S5).

We synthesized all 16 designs, of which 15 expressed. By biolayer interferometry (BLI), none of the 15 designs showed detectable binding against apo-*β*_1_AR in its inactive state, but 12 designs showed detectable binding when tested in the presence of the agonist isoprenaline, which stabilizes *β*_1_AR in an active state (Extended Data Fig. 5). The most potent binders NbIC6 and NbIC20 bind with higher affinities (13 and 21 nM, respectively) than Nb6B9 (26 nM) (Fig. 5c; Extended Data Fig. 5).

To evaluate design accuracy, we determined the cryo-electron microscopy (cryo-EM) structure of isoprenaline-bound *β*_1_AR in complex with NbIC20 (Fig. 5f; Extended Data Fig. 6). NbIC20 binds to the intracellular interface of *β*_1_AR and stabilizes the receptor in an active conformation, closely recapitulating the computational design model, with an RMSD of 0.84 Å over the nanobody alone and 0.76 Å over the active-state receptor alone. The CDR2 and CDR3 conformations are similar between the experimental structure and the design model, with RMSDs of 0.13 Å and 0.45 Å, respectively. CDR1 shows a modest local deviation from the design model (1.18 Å RMSD), but the overall binding mode and interface are well preserved (Fig. 5g). Together, these data demonstrate that NbIC20 binds *β*_1_AR as designed and that our model can design conformation-specific intracellular nanobodies against GPCRs with near-atomic structural accuracy.

To test whether GuideFlip produces more natural protein sequences than standard hallucination, we produced another 8 sequences in an ablation control, where we replaced the guided discrete flow with BindCraft’s soft and hard gradient descent over the CDRs, followed by SolubleMPNN^13^ redesign of the remaining CDR positions, with the interface residues and the framework held fixed (Supplementary Table S5). The hallucination-only designs had an average interface hydrophobicity of 48.3%, compared with 40.5% for the GuideFlip designs. The hallucination-only designs led to approximately four-fold lower amounts of expression than those from GuideFlip (Fig. 5d,e; Supplementary Table S6). However, probably because the intracellular binding site of the nanobody is relatively hydrophobic, the hallucination-only route did provide the most potent binder, NbIC19, with an affinity of *<*1 nM. These results suggest a potential trade-off between binding affinity and soluble expression at the hydrophobic intracellular surface of *β*_1_AR. As the use of the prior in GuideFlip is modular, we can use this result to trade soluble expression for higher affinity in cases with hydrophobic interfaces in our method.

## Discussion

The guided discrete flow matching approach in GuideFlip provides a general framework for coupling learned protein sequence distributions to structural objectives during generation. Rather than generating a backbone and subsequently finding a compatible sequence, the sequence and predicted structure evolve together. The sequence model constrains the search towards plausible proteins, whereas gradients from the structure-prediction model favour sequences satisfying the structural objective. The applications studied here span binding-induced folding of a disordered target, recognition of an intrinsically disordered region without a predefined bound structure and redesign of flexible loops. These problems can be addressed within the same framework by changing which residues are designable, which regions are structurally constrained and which interactions are favoured.

The sequence prior and structural guidance have distinct roles and can therefore be modified independently. In the present implementation, the sequence prior is switched on only after the binder reaches sufficient structural confidence; future approaches could instead vary its contribution continuously during generation. Here, an inverse-folding model trained on soluble proteins reduced the hydrophobic bias of direct AlphaFold optimization and improved nanobody expression. More generally, the prior could encode sequence properties appropriate to a particular design problem. Models trained on antibody repertoires, for example, could favour human-like antibody sequences, whereas family-specific evolutionary models could constrain enzyme design towards sequence distributions compatible with an existing fold or mechanism. Other sequence models could similarly introduce preferences related to solubility, stability or cellular context^44,45^.

The structural guidance can likewise be generalized. Although GuideFlip currently uses AlphaFold-derived objectives for target–binder complexes, the same principle could incorporate other differentiable predictors or scoring functions. Positive and negative objectives could favour one conformational state while disfavoring another, select between oligomeric assemblies or stabilize particular allosteric states. Objectives describing active-site geometry or catalytic residue placement could similarly broaden the approach beyond binder design. Its molecular scope could also extend beyond protein–protein interactions. ADFlip can represent atomic environments containing nucleic acids, small molecules and ions, and coupling such sequence priors to structural guidance from models that predict general biomolecular complexes, such as AlphaFold3^23^, could allow protein sequence and the conformations of interacting molecules to be explored together. This may be particularly valuable for RNA and flexible ligands, where function may depend on conformational changes in both partners.

More broadly, guided discrete flow matching provides a modular way to combine learned molecular priors with differentiable structural or functional objectives. This separation of prior and guidance offers a route towards joint sequence–structure design across molecular systems and functions.

## Methods

### ADFlip all-atom inverse folding

ADFlip^21^ is an all-atom inverse folding model that predicts amino-acid probabilities from structural and sequence context. Its multiscale graph neural network processes the spatial arrangement of residues through graph message passing and local atomic geometry through attention. Information is exchanged between the residue and atom representations before a Transformer decoder predicts amino-acid probabilities, conditioned on the partially specified sequence and time *t*.

For GuideFlip, we retrained ADFlip on a dataset of experimentally determined structures from the Protein Data Bank, excluding membrane proteins. During training, amino-acid identities are randomly masked according to a sampled time *t*, and side chains at masked positions are removed. The model learns to recover the original amino-acid identities from the remaining sequence and structural context using a cross-entropy loss.

During binder design, ADFlip receives the target and binder backbones, target side chains and partially specified binder sequence *s_t_*(Fig. 1b). Its output provides the sequence prior *p*_ADFlip_ at each binder position. The network parameters remain fixed throughout design. Details about the network architectures that underlie these steps and how they are trained have been described previously^21^.

### Backpropagation through AlphaFold

We use the differentiable AlphaFold-Multimer (v2.3) implementation in ColabDesign, following BindCraft^18^. AlphaFold receives the binder amino-acid probabilities, the target sequence and structural templates for selected regions. Target templates are retained for structured regions and omitted for flexible regions, allowing the latter to change conformation during complex prediction. The AlphaFold network parameters remain fixed throughout design.

The design loss, *L*_AF_, combines confidence and structural terms following BindCraft^18^. Confidence terms are derived from binder pLDDT, interface iPTM and predicted aligned errors within the binder and across the interface (PAE and iPAE). Structural terms describe predicted residue contacts within the binder and between the proteins, binder compactness measured by its radius of gyration, and helicity. The weights for the AlphaFold losses are provided in the Supplementary Table S7, with the helicity weight chosen according to the intended binder fold.

We backpropagate *L*_AF_ through AlphaFold to obtain gradients with respect to the binder input probabilities. These gradients are used to approximate the guidance factor *p*_AF_, favouring amino-acid changes predicted to reduce the design loss.

### GuideFlip sequence and structure codesign

Each GuideFlip update (Fig. 1b) takes the current amino-acid probabilities *p_t_* and complex *X_t_*. ADFlip predicts a sequence prior from the complex, which is combined with *p_t_*to form the binder input to AF-Multimer. AF-Multimer predicts the updated complex *X_t_*_+Δ*t*_, and backpropagation of the resulting design loss refines the combined amino-acid probabilities to give *p_t_*_+Δ*t*_.

The contribution of ADFlip is controlled by *λ_t_*. Initially, *λ_t_*= 0, and *p_t_*is updated solely using gradients of the AF-Multimer design loss. When the loss no longer improves and the predicted binder structure is sufficiently confident, we set *λ_t_*= 1. The ADFlip sequence prior then contributes alongside AF-Multimer guidance until the final binder sequence is obtained.

The design loss is calculated from the target–binder complex structure and confidence scores predicted by AF-Multimer. The updated amino-acid probabilities and the complex predicted before the gradient correction are passed to the next round.

Generation starts from random *p*_0_, with all designable binder positions remaining masked during AlphaFold-only updates. After ADFlip is enabled, i.e. when *λ_t_*= 1, we assign the most probable amino acid at positions where its probability reaches a decoding threshold. The probabilities continue to be updated until the binder sequence is fully specified. We estimate *t* from the reduction in sequence entropy. The sampling procedure is described in Algorithm 1 of the Supplementary Information.

### Design protocol and computational evaluation

The input to GuideFlip is the target protein structure, or its sequence alone when no structure is available; one can mark the flexible region of the target, the hotspot residues to target, the binder length, and the number of design trajectories to run.

Structure templates may be used to control which parts of the complex are free to move. Structured regions of the target are supplied as a template and kept close to their input conformation. We remove the template for flexible regions so that they can refold against the binder at every step; a fully disordered target is run with no template at all. For nanobody design the same control is applied on the binder side: the VHH framework is templated and held fixed, and only the CDR loops are left free to be designed.

After design, we evaluated each target–binder complex with AlphaFold3 (version 3.0.0)^23^. Binder sequences were supplied without homologous multiple sequence alignment (MSA) information. Binder templates were omitted except for nanobodies, for which only VHH framework residues were included and CDR regions were excluded. For nanobody designs, we ran AlphaFold3 with 20 random seeds and retained the highest-scoring model as the predicted complex.

Target MSAs were retained for RBX1 and *β*_1_AR, whereas cropped *α*-synuclein was evaluated using only its query sequence and without structural templates. Where target templates were used, flexible regions were excluded from the template mappings, allowing these regions to adopt a bound conformation during prediction.

Designs were assessed using interface confidence (iPTM), binder confidence (pLDDT), and minimum interface predicted aligned error (iPAE_min_). Agreement between the design model and the AlphaFold3 prediction was assessed using interface root-mean-square deviation (iRMSD), calculated with the DockQ package^46^ (Fig. 2). Filtering thresholds were set separately for each target. For nanobodies, binder pLDDT was evaluated over the CDR loops only.

For fragment-based DisProt designs, the AlphaFold3 prediction for the full-length target was cropped to the corresponding target fragment while retaining the binder. Interface RMSD was then calculated with DockQ 2.1.3 by superposing the corresponding interface residues from both chains.

We assessed robustness using four ADFlip–AlphaFold codesign runs started from the same lowest-loss state retained during the initial stages of AlphaFold-only optimization. The AlphaFold gradients and entropy-derived time variable can differ between runs, producing alternative binder sequences. Each sequence was evaluated with AlphaFold3 using the same confidence and pose-agreement criteria. A robustness score was defined as the fraction of the four sequences that pass all criteria; a score of 1 indicates that all four pass. Designs were ranked by this score.

### Expression and biotinylation of α-synuclein

A construct encoding wild-type *α*-synuclein was cloned into pRK172. The plasmid was transformed into BL21(DE3)-Gold competent cells (Agilent), which were grown on a 2*×*TY agar plate containing ampicillin. The transformed cells were used to inoculate 0.5–2 L of 2*×*TY medium supplemented with 2.5 mM magnesium sulfate and 100 mg/L ampicillin. The cells were cultured at 37 *^◦^*C until the OD_600_ reached 0.6. Isopropyl *β* -D-1-thiogalactopyranoside (IPTG) was then added to a final concentration of 1 mM to induce expression. After induction for 3–4 h at 37 *^◦^*C, the cells were pelleted by centrifugation at 2,500 g for 20 min at 4 *^◦^*C. The pellets were flash-frozen in liquid nitrogen and stored at *−*80 *^◦^*C.

Frozen cell pellets were resuspended in 50 mL of cold lysis buffer containing 10 mM Tris (pH 7.4), 5 mM EDTA and one protease inhibitor cocktail tablet. The cells were sonicated at 40% amplitude for a total sonication time of 2 min, in cycles of 30 s on and 30 s off. The lysate was centrifuged at 18,000 rpm for 45 min at 4 *^◦^*C to remove large debris. The pH of the supernatant was reduced to 3.5, and the solution was stirred for at least 20 min at room temperature before being centrifuged at 50,000g for 1 h. The pH of the resulting supernatant was then adjusted to 7.4 with sodium hydroxide. The protein was precipitated by adding ammonium sulfate to a final concentration of 0.33 g/mL and pelleted by centrifugation at 18,000 rpm for 30 min at 4 *^◦^*C. The pellet was dissolved in cold PBS and purified further on a HiLoad 16/600 Superdex 75 pg column (GE Healthcare) at 1 mL/min. Pure fractions were identified by sodium dodecyl sulfate–polyacrylamide gel electrophoresis (SDS–PAGE), pooled and concentrated to approximately 6 mg/mL in a Vivaspin concentrator with a 3 kDa molecular-weight cut-off (Sartorius) before being aliquoted.

EZ-Link Sulfo-NHS-Biotin (2 mg; Thermo Fisher Scientific) was dissolved in 600 *µ*L of dimethyl sulfoxide. The *α*-synuclein monomer was centrifuged at 55,000 rpm for 1 h at 4 *^◦^*C to remove any potential aggregates. The *α*-synuclein solution was mixed with biotin at a ratio of 1:20 and left to react for 30 min at room temperature. Excess biotin was removed on a Zeba Spin Desalting Column (Thermo Fisher Scientific). Biotinylation was confirmed by western blotting, and the biotinylated protein was used for SPR.

^15^N-labelled *α*-synuclein was expressed in M9 medium containing ^15^N-labelled ammonium chloride as the sole nitrogen source. Expression was induced overnight with 1 mM IPTG. The labelled protein was purified as described above and concentrated to approximately 20 mg/mL for NMR.

### Expression of α-synuclein binders

Binder genes were cloned into the pET-3a vector and transformed into BL21(DE3)-Gold competent cells. The transformed cells were used to inoculate 1–2 L of 2*×*TY medium supplemented with 100 mg/L ampicillin. The cells were cultured at 37 *^◦^*C until the OD_600_ reached 0.6. IPTG was then added to a final concentration of 1 mM, and expression was induced overnight at 18 *^◦^*C and 220 rpm. The pellets were resuspended and sonicated, and the clarified supernatant was incubated with Ni–NTA resin. The resin was washed first with 20 mM HEPES (pH 7.2), 500 mM NaCl, 10 mM imidazole and then with 20 mM HEPES (pH 7.2), 100 mM NaCl, 40 mM imidazole. The bound protein was eluted with 300 mM imidazole and exchanged into 20 mM HEPES (pH 7.2), 100 mM NaCl by dialysis or on a Zeba Spin Desalting Column (Thermo Fisher Scientific) before use in SPR. For NMR, the binders were purified further on a HiLoad 16/600 Superdex 75 pg column (GE Healthcare) in 20 mM HEPES (pH 7.2), 100 mM NaCl after Ni–NTA affinity purification, and the pure peak fractions were pooled and concentrated.

### Surface plasmon resonance (SPR) of α-synuclein binders

SPR was performed on a Biacore T200 using SA sensor chips (Cytiva). Both the reference and the analyte channels were equilibrated in 20 mM HEPES (pH 7.2), 50 mM NaCl, 0.05% Tween-20 at 20 *^◦^*C. Biotinylated *α*-synuclein was immobilized on the chip surface to approximately 486 RU. Designed binders and control antibodies were first screened at a single concentration of 200 *µ*M. The affinities of the positive binders were then measured using concentration series as indicated in the individual figures. In both experiments the analyte was injected over the chip for 120 s at 30 *µ*L/min, with a 900 s dissociation time. The surface was regenerated with a 60 s injection of 2 M NaCl at 30 *µ*L/min. The data were double-referenced against a blank but similarly modified flow channel, and a buffer-only injection was subtracted. After reference and buffer correction, the sensorgrams were fitted in Prism 11.0.2 (GraphPad Software). The responses at equilibrium (*R*_eq_) were fitted to a 1:1 binding model with a constant background term to give *K*_D_:

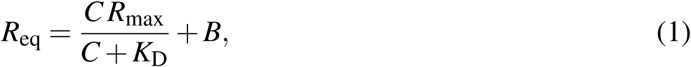

where *C* is the analyte concentration, *R*_max_ is the maximum response at saturation and *B* is the background resonance.

### Nuclear magnetic resonance (NMR) of α-synuclein binders

NMR data were collected at 15 *^◦^*C on a Bruker Avance III HD spectrometer operating at a proton frequency of 700 MHz and fitted with a 5 mm TXO cryoprobe optimized for carbon and nitrogen sensitivity. Samples were prepared in 20 mM HEPES (pH 7.2), 50 mM NaCl and 10% (v/v) D_2_O as the lock solvent. In these experiments, the ^15^N-labelled *α*-synuclein was held at 100 *µ*M and the amount of dnAS15 or dnAS36 was varied.

The interaction of the binders with ^15^N-labelled *α*-synuclein was confirmed by changes in absolute peak intensity in the ^15^N HSQC (fHSQC, Bruker sequence library), with peak assignments taken from BMRB entry 52093^47^ and BMRB entry 19337^48^. Small chemical shift perturbations (CSPs) were quantified as in equation (2); they could be attributed to minor changes in pH over the course of the titration^49^:

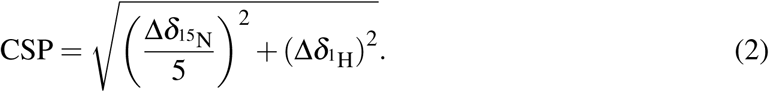

Transverse relaxation rates *R*_2_ were measured with pseudo-3D experiments (hsqct2etf3gpsi3d, Bruker pulse sequence library) and a recovery delay of 3 s. Relaxation delays of 16.96 (*×*2), 33.92, 50.88, 67.84, 84.8, 101.7, 135.68, 152.64, 169.6, 203.52 and 254.4 ms were acquired in an interleaved fashion, and the rates were obtained by fitting the intensities to an exponential decay. All data were processed in Bruker TopSpin 3.6.5 or 4.5.0 and analysed in POKY^50^.

### RBX1 protein production

Recombinant human RBX1 (UniProt P62877) was produced by Data Powered Therapeutics GmbH (DPTx). Two constructs were made: full-length RBX1 (residues 1–108) and an N-terminally truncated construct (residues 36–108). Both were expressed as N-terminal 8*×*His–MBP fusions to improve soluble yield. The seven competition designs were tested against the full-length construct. Protein was purified by immobilized metal-affinity chromatography on a Ni^2+^ resin, and the fusion tag was removed by on-column cleavage with human rhinovirus 3C protease, releasing untagged RBX1. The cleaved protein was purified further by size-exclusion chromatography, concentrated, flash-frozen in liquid nitrogen and stored at *−*80 *^◦^*C until use. Protein quality and monodispersity were assessed by analytical size-exclusion chromatography and flow-induced dispersion analysis. The final protein was formulated in 20 mM HEPES (pH 7.5) and 150 mM NaCl.

### Cloning and expression of RBX1 binders

Binders were expressed and tested by Adaptyv Bio. Coding sequences were reverse-translated from the designed protein sequences, codon-optimized, and ordered as gene fragments (Twist Bioscience) together with a 3*^′^* fragment carrying a linker and a C-terminal Twin-Strep tag. Fragments were assembled with the NEBuilder HiFi DNA Assembly kit (New England Biolabs) in 2 *µ*L reactions, checked by capillary electrophoresis (Agilent ZAG DNA Analyzer) and quantified with a Qubit assay (Invitrogen). Each binder was expressed in an 8 *µ*L reaction using an optimized prokaryotic *in vitro* translation system and 4 nM assembled gene fragment, incubated at 37 *^◦^*C for 8 h. Concentration and yield were then normalized by affinity-based quantification. The binders were not purified: each tagged binder was captured directly from the crude cell-free reaction onto the biosensor through its Twin-Strep tag, so every measurement was made on crude material with tag-specific, on-sensor capture.

### SPR of RBX1 binders

The seven competition designs (Round 1) were measured in triplicate by SPR on a Carterra LSA XT instrument. Twin-Strep-tagged binders were captured on a carboxymethylated sensor chip functionalized with Strep-Tactin XT (IBA Lifesciences; 50 *µ*g/mL in 10 mM sodium acetate, pH 4.5) by EDC/NHS coupling, and the surface was quenched with 1 M ethanolamine hydrochloride. The running buffer contained 10 mM HEPES (pH 7.4), 150 mM NaCl, 3 mM EDTA and 0.05% Tween-20. RBX1 was injected as the analyte at 10.0, 31.6, 100, 316 and 1000 nM, with 300 s association and 600 s dissociation at a flow rate of 50 *µ*L/min, and the surface was regenerated with 10 mM glycine-HCl (pH 1.5). Binding constants were extracted by globally fitting a 1:1 Langmuir model to the reference-subtracted data using Adaptyv fitting software.

### RBX1–dnRB1 interface mutagenesis

Binder residues that buried solvent-accessible surface area on complex formation (ΔSASA *>* 10 Å^2^) were grouped by C*α* position into five spatial clusters (A–E; Fig. 4d). For the single-cluster mutants, residues were substituted with alanine, except in charged cluster B, which was tested by charge reversal. Single-cluster and combined-cluster mutants were assessed for RBX1 binding by BLI as described below.

### Biolayer interferometry (BLI) of RBX1 binders

The five second-round designs (Round 2) and the dnRB1 mutation panel were measured in triplicate by BLI on a Gator Bio instrument, using Strep-Tactin XT probes to capture the Twin-Strep-tagged binders. The running buffer contained 10 mM HEPES (pH 7.4), 150 mM NaCl, 3 mM EDTA and 0.2% Tween-20. Probes were equilibrated in running buffer, and each binder was loaded for 120 s to a target shift of 0.5–1.0 nm, between a 120 s first baseline and a 200 s second baseline. RBX1 was then injected as the analyte in a half-log dilution series from 1000 to 30 nM (four concentrations), with 220 s association and 240 s dissociation. Probes were regenerated between cycles in 10 mM glycine-HCl (pH 1.5), applied five times for 10 s, and neutralized in running buffer. Buffer-only and non-binding-binder controls were included for reference subtraction and drift correction. Measurements were made at 25 *^◦^*C with data sampled at 5 Hz. Sensorgrams were trimmed to the baseline, association and dissociation phases, corrected for signal jumps at the phase transitions, aligned, and baseline- and reference-subtracted. The binding constants were extracted by globally fitting a 1:1 Langmuir model across all concentrations using Adaptyv fitting software; in the global fit *k*_off_ and *K*_D_ were fitted directly and *k*_on_ was calculated as *k*_off_*/K*_D_. Designs that gave an association-phase signal at least 300% above the negative control but could not be fitted reliably were considered binders, without a reported *K*_D_. Binding outcomes for dnRB1 interface mutants were summarized using the assay-provider classifications. Relative to wild-type dnRB1, constructs classified as strong or medium binders were labelled as retained or weakened binding, respectively. Constructs reported as non-binding were labelled as no detectable binding under the assay conditions.

### Expression and biotinylation of β_1_AR

A construct encoding *β*_1_AR with an N-terminal 2x AviTag and a C-terminal 8xHis tag (2xAviTag-*β*_1_AR-His8) was cloned into pAcGP67B (BD Biosciences). High-titre recombinant baculoviruses were prepared using the flashBAC ULTRA system (Oxford Expression Technologies). *Trichoplusia ni* High Five cells (Thermo Fisher Scientific) were grown in suspension in ESF 921 medium (Expression Systems). Insect cells at 2 *×* 10^6^ cells/mL were infected with 2% (v/v) *β*_1_AR baculovirus and cultured for 48 h. The cells were harvested by centrifugation, flash-frozen in liquid nitrogen, and stored at *−*80 *^◦^*C until further use. The membrane fraction was prepared through two rounds of homogenization using an Ultra-Turrax disperser (IKA) and centrifugation at 40,000 g for 1 h in 1x PBS, 350 mM NaCl. The membranes were solubilized in 1.7% (w/v) lauryl maltose neopentyl glycol (LMNG, Anatrace) and further purified with 0.005% LMNG by Ni^2+^-affinity chromatography as previously described^51^. The eluate was concentrated using a 100-kDa molecular weight cut-off (MWCO) Amicon Ultra centrifugal concentrator (Merck) and passed over a Superdex 200 Increase column pre-equilibrated in 1x PBS, 100 mM NaCl, 0.005% (w/v) LMNG, and 0.2 mM EDTA. The peak fractions containing *β*_1_AR were pooled and concentrated. Purified *β*_1_AR was enzymatically biotinylated using BirA biotin ligase (Avidity) overnight. The reaction mixture was subsequently desalted to remove excess free biotin, and biotinylated *β*_1_AR was eluted in 1*×* PBS, 100 mM NaCl, 0.2 mM EDTA and 0.005% (w/v) LMNG.

### Expression of the nanobodies against β_1_AR

Synthetic genes (Integrated DNA Technologies) for designed nanobodies were cloned into plasmid pET-29b (Novagen) with an N-terminal 6xHis tag followed by a thrombin protease cleavage site. Expression in *Escherichia coli* (*E. coli*) strain BL21(DE3) RIL (Agilent Technologies) and purification from the periplasmic fraction were as described elsewhere^41^.

### BLI of β_1_AR binders

The binding kinetics of *β*_1_AR and nanobodies were measured by BLI on an Octet RED384 instrument (ForteBio). The experiments were conducted at 30 *^◦^*C. Biotinylated *β*_1_AR was diluted in buffer containing 20 mM HEPES (pH 7.4), 150 mM KCl, 10 mM MgCl2, 0.005% LMNG and 10 *µ*M isoprenaline, and immobilized onto Octet Streptavidin (SA) Biosensors (Sartorius). After washing twice in the same buffer, both association and dissociation were measured with the nanobody in different concentrations. The binding rate constants were extracted by globally fitting a 1:1 Langmuir model to the data using Octet Data Analysis HT software (v11.1).

### Formation of isoprenaline-bound β_1_AR-NbIC20 complex

Preparation of the receptor-nanobody complex was performed as described previously^38^. *β*_1_AR (0.8 mg) was mixed with a 2.5-fold molar excess of nanobody (1 mg), and isoprenaline was added to a concentration of 10 *µ*M in a final volume of 100 *µ*L. The mixture was then incubated for 2 h on ice. After incubation, size-exclusion chromatography (SEC) was performed to separate the receptor-nanobody complex from excess nanobody. The complex was loaded and separated on a Superdex 200 10/300 GL column (GE Healthcare), pre-equilibrated with 20 mM HEPES (pH 7.4), 150 mM KCl, 10 mM MgCl2, and 0.005% LMNG. The peak fractions corresponding to the complex were concentrated to a final concentration of 10 mg/mL.

### β_1_AR-NbIC20 cryo-electron microscopy

Cryo-EM grids were prepared by applying 3 *µ*L of the sample (at a protein concentration of 10 mg/mL) on plasma-treated holey gold grids (UltrAufoil Au 1.2/1.3 300 mesh). Excess sample was removed by blotting with filter paper for 4 s before plunge-freezing in liquid ethane (cooled to *−*181 *^◦^*C) using an FEI Vitrobot Mark IV maintained at 100% relative humidity and 4 *^◦^*C. Images were collected on an FEI Titan Krios microscope at 300 kV on a Falcon 4i electron detector using EPU software 3.8 (Thermo Fisher Scientific). Micrographs were collected as movies at approximately 1.0 electron/Å^2^/frame, with a total fluence of 60 electrons/Å^2^, at a magnification of 120,000*×* (0.67 Å/pixel).

### β_1_AR-NbIC20 cryo-EM image processing

Cryo-EM images were processed in RELION 5.0^52^. Overall, drift, beam-induced motion and dose weighting were corrected using RELION’s own implementation of MotionCor2^53^. Contrast transfer function (CTF) fitting and phase shift estimation were performed using CTFFIND4.1^54^. Particles were picked automatically using Laplacian-of-Gaussian, extracted in a box size of 320 pixels and downsampled four times, and subjected to two rounds of 2D classification, one round of 3D classification, and another round of 2D classification to select a clean subset. Selected particles were re-extracted with a box size of 360 pixels, without downsampling, for 3D auto-refinement with Blush regularization^55^, yielding high-resolution density maps with a global resolution of 3.3 Å. The resolution of the final map was estimated and the map was sharpened using standard post-processing procedures in RELION.

### *β*_1_*AR-NbIC20 cryo-EM atomic modelling*

The initial *β*_1_AR-NbIC20 model was generated using ModelAngelo^56^. Ligand CIF files were generated using AceDRG^57^. After docking the models into the corresponding EM density maps using UCSF ChimeraX^58^, iterative rounds of manual adjustment and real-space refinement were performed in Coot^59^ and Phenix^60^, respectively. MolProbity^61^ was used to validate the final models, and the comprehensive refinement statistics are provided in Extended Data Table 1.

## Data availability

Data supporting the findings of this study are available in the main text, Supplementary Information and the resources listed below. The cryo-EM map of the *β*_1_AR–NbIC20 complex has been deposited in the Electron Microscopy Data Bank (EMDB) under accession code EMD-59993. The corresponding atomic coordinates have been deposited in the Protein Data Bank (PDB) under accession code 34KK. The database of computationally designed binders is available at https://huggingface.co/spaces/ykiiiiii/guideflip-idp-binder.

## Code availability

GuideFlip is freely available under the MIT licence at https://github.com/ykiiiiii/Guid eFlip.

## Acknowledgements

We thank I. Clayson, T. Darling and J. Grimmett for help with high-performance computing, and the EM facility of the Medical Research Council (MRC) Laboratory of Molecular Biology for help with cryo-EM data acquisition. We thank Keren Turton of the Tissue Culture Facility at the MRC Laboratory of Molecular Biology for assistance with insect cell culture. We also thank Data Powered Therapeutics GmbH (DPTx) for producing the recombinant RBX1 target protein, and Adaptyv Bio for expressing and testing the RBX1 binders in both design rounds. The authors acknowledge the use of resources provided by the Isambard-AI National AI Research Resource (AIRR). Isambard-AI is operated by the University of Bristol and is funded by the UK Government’s Department for Science, Innovation and Technology (DSIT) via UK Research and Innovation and the Science and Technology Facilities Council [ST/AIRR/I-A-I/1023].

## Author contributions

K.Y. designed and implemented GuideFlip, with conceptual contributions by K.J. and P.T.; Q.C. and C.G.T. performed experimental validation of *β*_1_AR binders and cryo-EM structure determination; D.Z. performed experimental validation of *α*-synuclein binders, with contributions to SPR by S.M. and to NMR by J.W.; K.J. and S.H.W.S. supervised the project. All authors contributed to the writing of the manuscript.

## Additional information

Correspondence and requests for materials should be addressed to K.Y., K.J. and S.H.W.S.

For the purpose of open access, the MRC Laboratory of Molecular Biology has applied a CC-BY public copyright license to any Author Accepted Manuscript version arising.

## Extended Data

**Extended Data Fig. 1:**
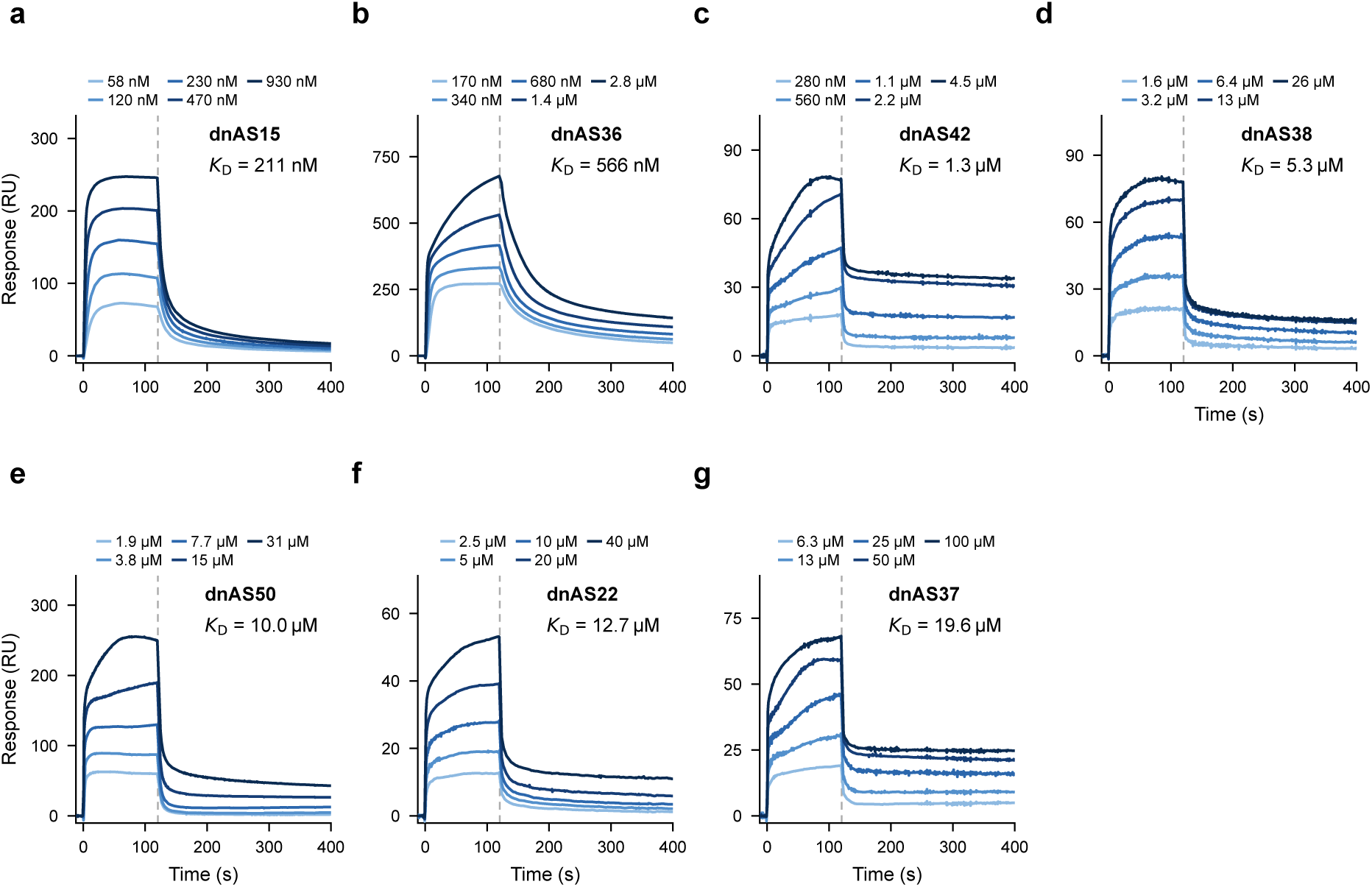
SPR sensorgrams of the seven *α*-synuclein binders. **a**–**g**, Sensorgrams for the seven designs with quantifiable binding, ordered from highest to lowest affinity. Five concentrations are shown for each binder, with darker blue indicating higher concentrations. Dashed vertical lines mark the onset of dissociation. Reported *K*_D_ values are indicated in each panel.

**Extended Data Fig. 2:**
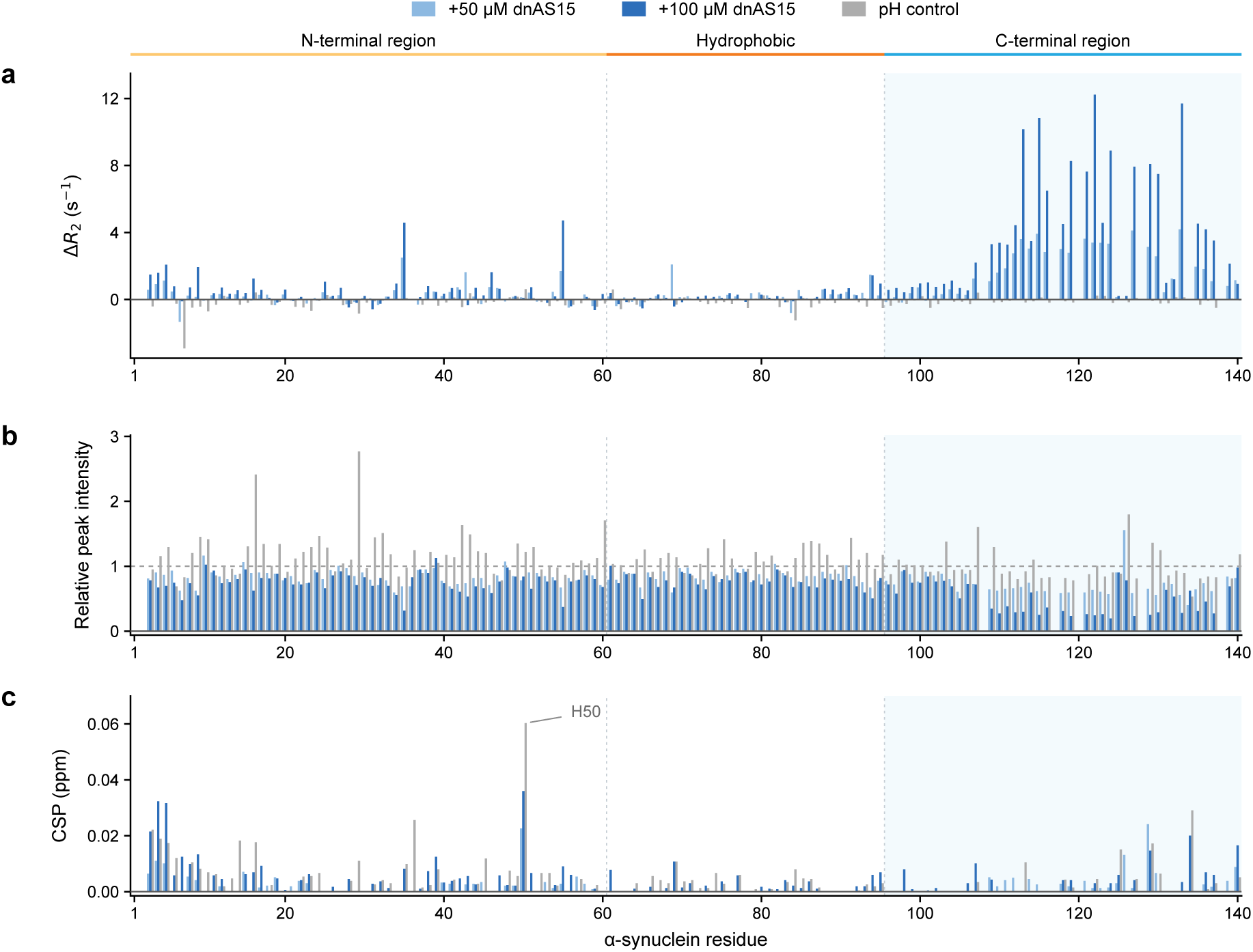
NMR characterization of *α*-synuclein binding to dnAS15. **a**, Per-residue changes in backbone transverse relaxation rates (Δ*R*_2_) for 100 *µ*M full-length *α*-synuclein with 50 or 100 *µ*M dnAS15, relative to *α*-synuclein alone, for 131 assigned backbone amides. **b,c**, Relative peak intensities (**b**) and chemical shift perturbations (CSP; **c**) for the indicated conditions. Light and dark blue denote 50 and 100 *µ*M dnAS15, respectively; grey denotes the pH control (*α*-synuclein with 1 *µ*l of 1 M HCl). Bars show point estimates. The amino-terminal (1–60), hydrophobic (61–95) and carboxy-terminal (96–140) regions are indicated above **a** and aligned across **a**–**c**; the carboxy-terminal region is shaded. The dashed line in **b** marks a relative intensity of 1. H50 is labelled in **c**.

**Extended Data Fig. 3:**
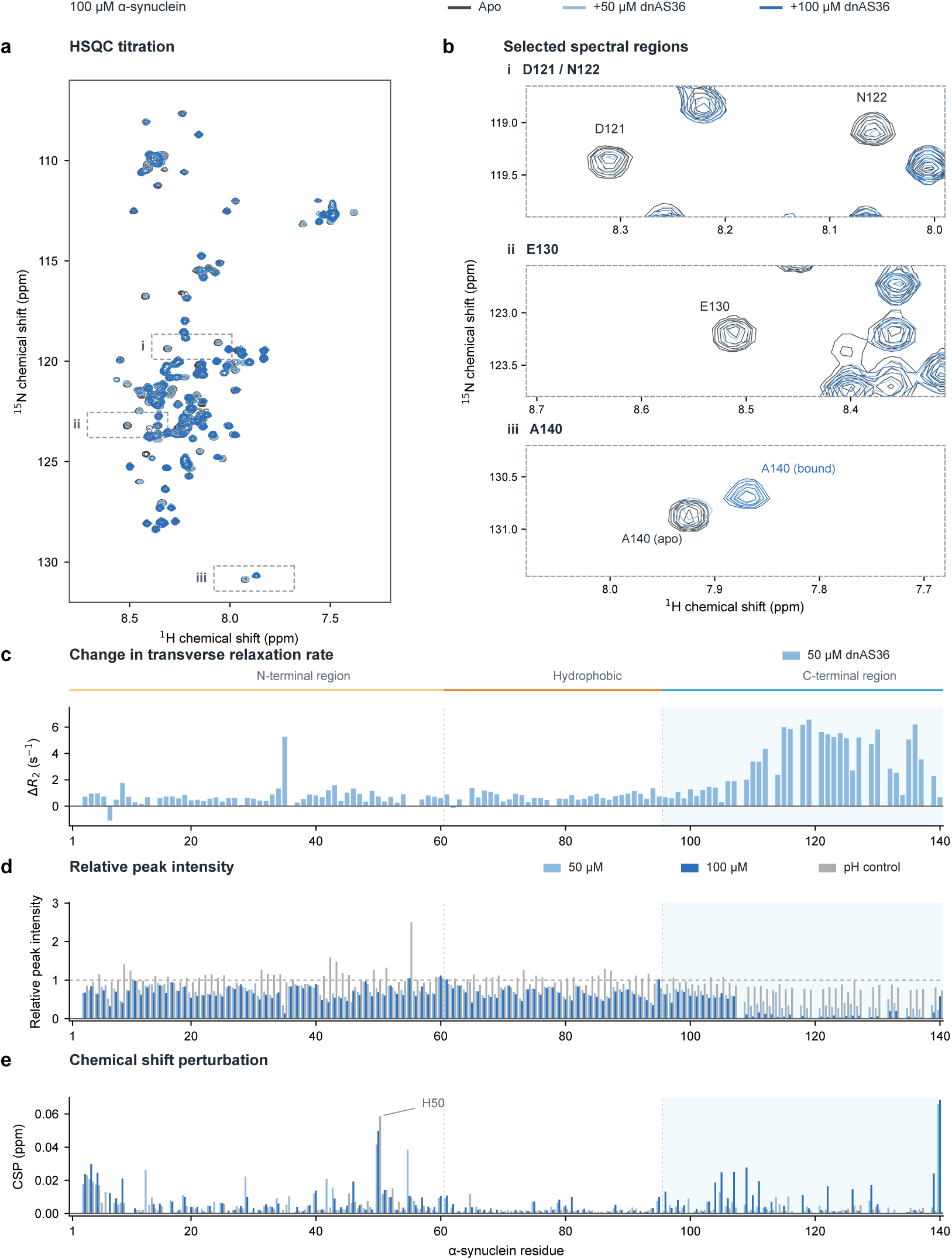
NMR analysis of dnAS36 binding to *α*-synuclein. **a**, ^1^H–^15^N HSQC spectra of 100 *µ*M *α*-synuclein alone (dark grey) or with 50 *µ*M (light blue) or 100 *µ*M (blue) dnAS36. **b**, Enlargements of the boxed regions in **a**. **c**, Per-residue changes in transverse relaxation rates (Δ*R*_2_) with 50 *µ*M dnAS36 relative to apo *α*-synuclein. **d,e**, Relative peak intensities (*I/I*_0_; **d**) and chemical shift perturbations (CSP; **e**) with 50 or 100 *µ*M dnAS36; grey bars indicate the pH control. Peak heights are normalized to their respective apo references. The bound-state A140 peak is used in **d,e**. Shading marks residues 96–140; the dashed line in **d** indicates *I/I*_0_ = 1.

**Extended Data Fig. 4:**
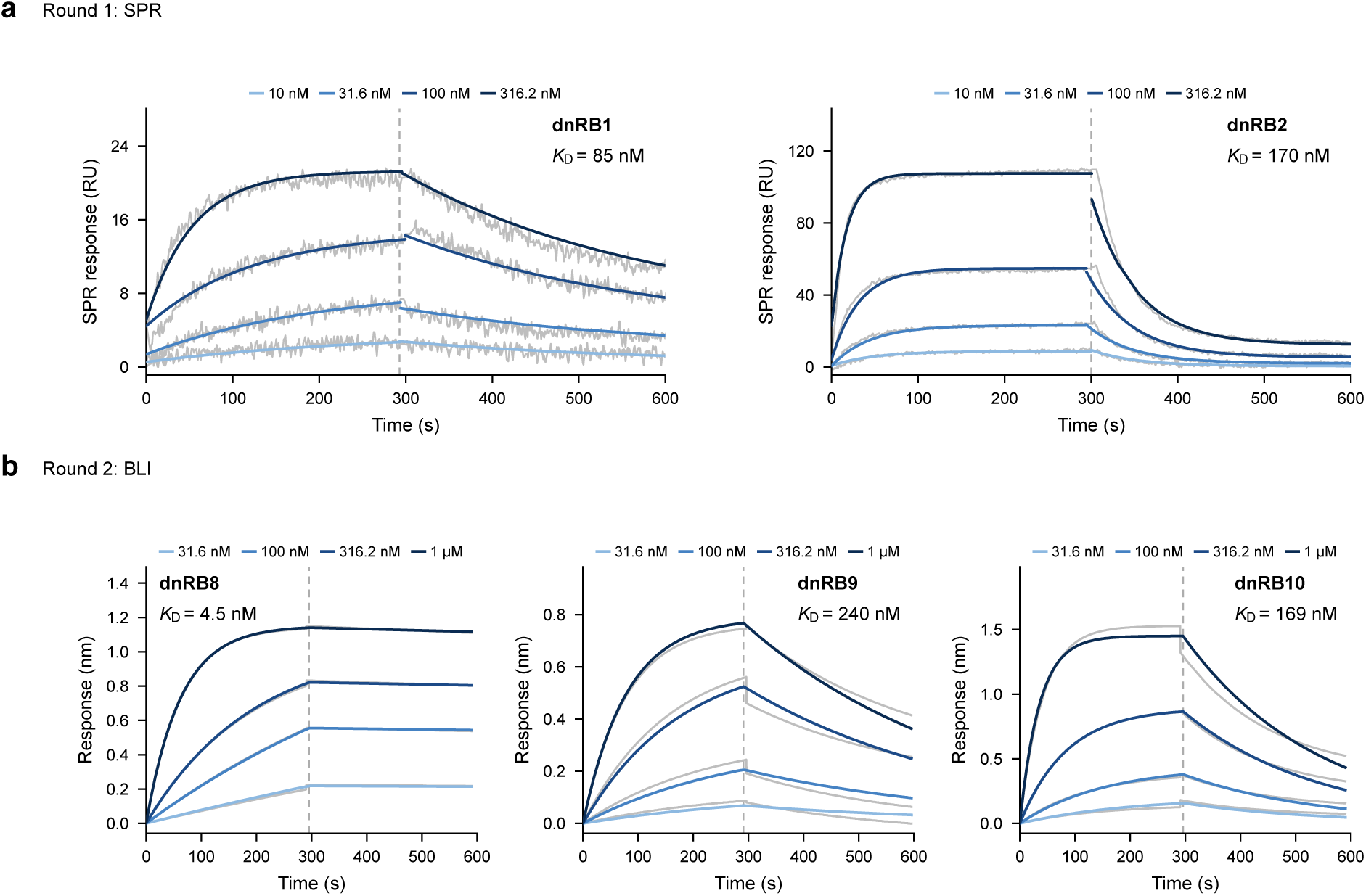
Binding kinetics of designed RBX1 binders. **a**, Surface plasmon resonance (SPR) sensorgrams for dnRB1 and dnRB2 from the first design round. **b**, Biolayer interferometry (BLI) sensorgrams for dnRB8, dnRB9 and dnRB10 from the second design round. Grey traces show measured responses; blue curves show global fits to a 1:1 binding model. Darker blue indicates higher concentrations. Concentrations and reported *K*_D_ values are indicated in each panel. Dashed lines mark the start of dissociation. The 1 *µ*M SPR traces were omitted from the display because they showed a hook effect. For dnRB8, the measurement with the higher *R*^2^ among the two available datasets is shown.

**Extended Data Fig. 5:**
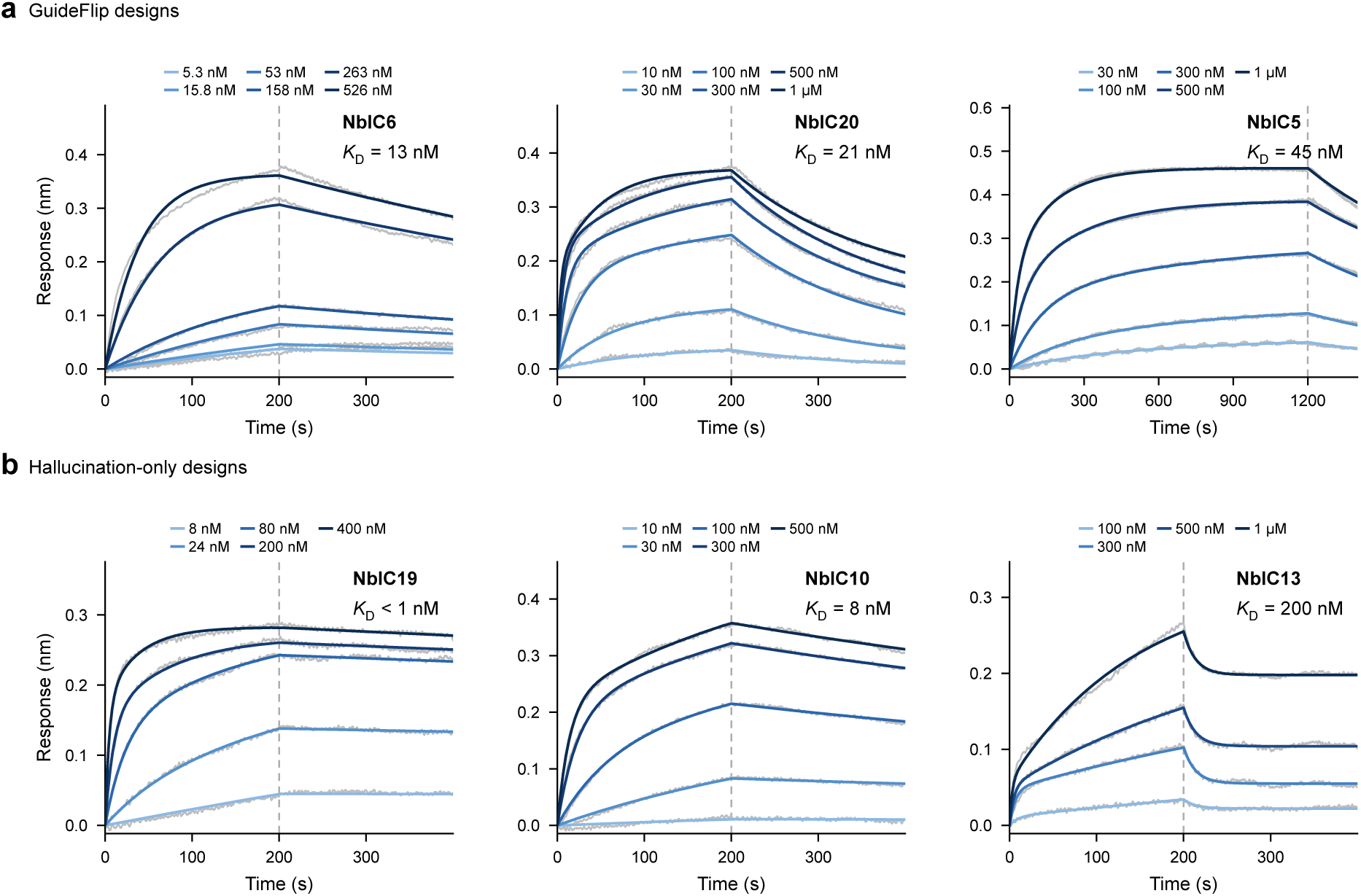
Binding kinetics of designed intracellular nanobodies against *β*_1_AR. **a,b**, Biolayer interferometry (BLI) sensorgrams for GuideFlip designs (NbIC6, NbIC20 and NbIC5; **a**) and hallucination-only designs (NbIC19, NbIC10 and NbIC13; **b**) binding to isoprenaline-bound *β*_1_AR. Grey traces show measured responses; blue curves show global fits to a 1:1 Langmuir binding model. Darker blue indicates higher nanobody concentrations. Concentrations and *K*_D_ values are indicated in each panel. Dashed lines mark the start of dissociation.

**Extended Data Fig. 6:**
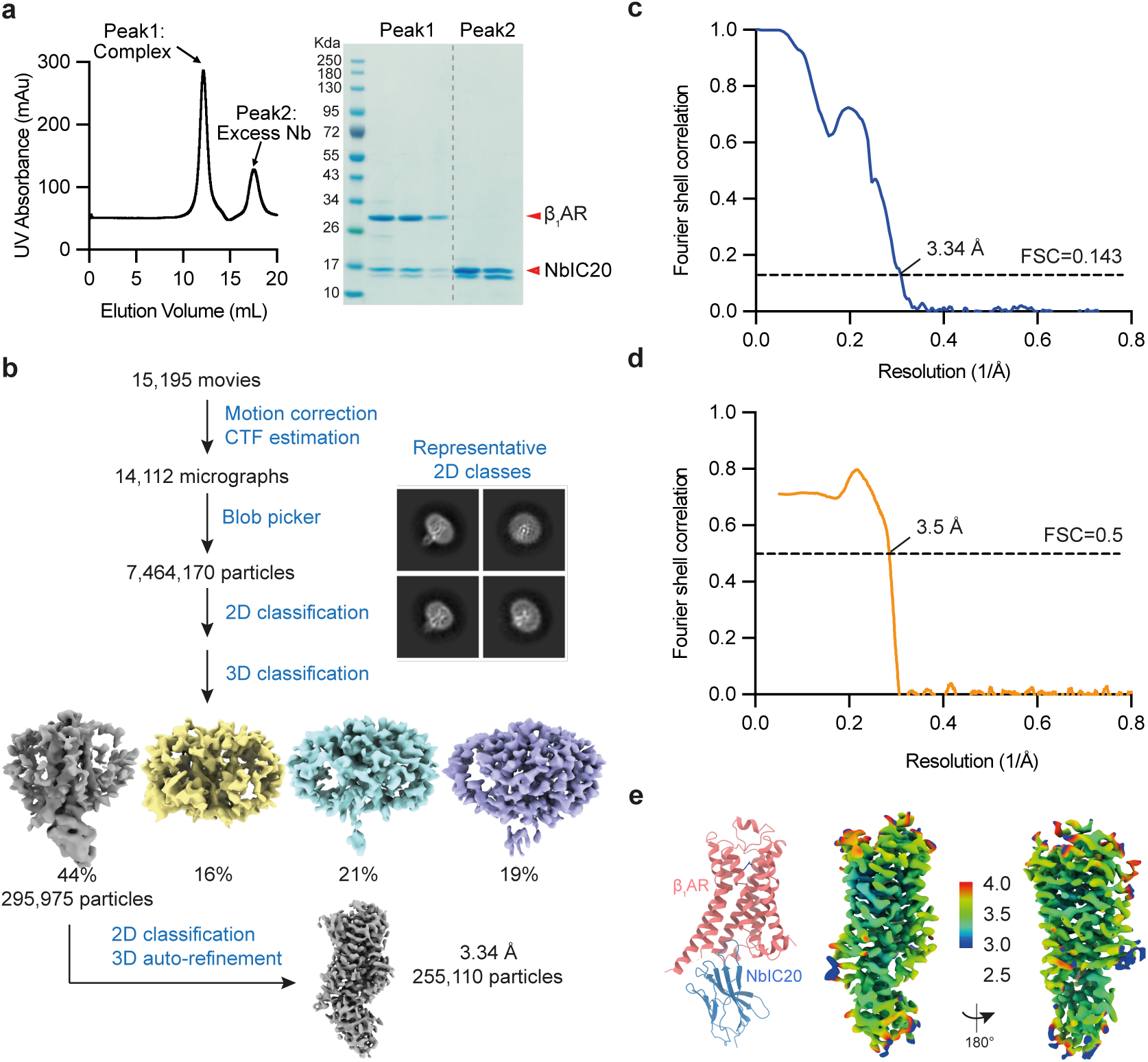
Biochemical characterization and cryo-EM analysis of the *β*_1_AR–NbIC20 complex. **a**, Size-exclusion chromatography profile (left) and SDS–PAGE analysis of fractions from the indicated peaks (right). Peak 1 contains the isoprenaline-bound *β*_1_AR–NbIC20 complex; peak 2 contains excess NbIC20. **b**, Cryo-EM data processing workflow, with representative two-dimensional class averages and three-dimensional classes. **c**, Gold-standard Fourier shell correlation (FSC) curve, indicating a map resolution of 3.34 Å at an FSC threshold of 0.143. **d**, Model–map FSC curve, crossing the FSC threshold of 0.5 at 3.5 Å. **e**, Atomic model (left) and cryo-EM map coloured by local resolution in two views related by a 180*^◦^* rotation. The colour scale indicates local resolution in Å.

**Extended Data Table 1:** Cryo-EM data collection, refinement and validation statistics for the isoprenaline-bound *β*_1_AR–NbIC20 complex.

| | $\beta_1$AR–NbIC20 |
| --- | --- |
| Magnification | 120,000x |
| Voltage (kV) | 300 |
| Electron exposure (e<sup>−</sup>/Å<sup>2</sup>) | 60 |
| Defocus range (μm) | −0.8 to −1.8 |
| Pixel size (Å) | 0.67 |
| Symmetry imposed | – |
| Map resolution (Å) | 3.34 |
| FSC threshold | 0.143 |
| **Refinement** |  |
| *Model composition* |  |
| Non-hydrogen atoms | 3137 |
| Protein residues | 394 |
| *R.m.s. deviations* |  |
| Bond lengths (Å) | 0.002 |
| Bond angles (°) | 0.827 |
| *Validation* |  |
| MolProbity score | 1.38 |
| Clash score | 3.33 |
| Poor rotamers (%) | 0 |
| *Ramachandran plot* |  |
| Favoured (%) | 96.91 |
| Allowed (%) | 3.09 |
| Disallowed (%) | 0 |

## Footnotes

1 https://huggingface.co/spaces/ykiiiiii/guideflip-idp-binder

