## Supplementary information for "De novo design of flexible protein interactions with GuideFlip"

Supplementary Table S1: **Amino-acid sequences of the 52 primary $\alpha$-synuclein designs.** Sequences are listed in blocks of ten residues, excluding affinity tags and associated cloning linkers. Design identifiers correspond to Supplementary Table S2.

| Design | Sequence |
|---|---|
| dnAS1 | MVTMEITPDK IVMHMPKEKM EKILKEMIPE VAGKMEEKGG MLTDEERDEM<br>TDKYVKTFQK KTGYKDEEEV YKWVRSMMKW VGKKIKERWM TG |
| dnAS2 | MASIEKTPEK ITLYIPREVF EEFMREFIPE LAARMEERGG PLSEEELEEL<br>AEKYVERLME LTGLKDREAV YEWVRSLCRL VQAKLLRYRE EL |
| dnAS3 | MKMEITPESI TLYVPRELFE KFMRELIPEI YERMKEKGGP LSEEEKEELA<br>EKYVERLMEL TGFEDREAVY KWVRSLIYLV QKKLLEYEKT L |
| dnAS4 | EQRAVIEMMT AMFRTYVGMP MTQEEFDELY HTSMRYMYWT QMDDEHFLEM<br>MRTFADIVKG HWKREQAATP NGAIDPGVQK TIWMLAGFVR HHTMWMIDKN ENPDM |
| dnAS5 | MQDEVREELL AEVRELVDKP VSDEEFREFW DRTMRRIVEA KMDRESLLEL<br>LRGMAEVVKG TWAKLKAENP DGKIDPGVKR TIYKLAAFVR HLTERLRRKE ENPEE |
| dnAS6 | EQEEVVEEIV KEVEELVTKP VSREEFNEFW EKLRKKIVET KMDRETLIEL<br>LRRMAEIVKG TWEALKAANP DGKIDPGVKR TIYMLAELVR HETERLRYKE ENPEL |
| dnAS7 | FQTYMMYKMD RMMTDMRKTA RVIIFLSDTY PDEEIIGDQG YEVKQQWPTA<br>RRQMREIEEE YYEIRTYGAR SMSEMRGLYY RMREMYRKIW RMAKKLIMGF<br>DWAKHPETID EVRYLLSVTT EPT |
| dnAS8 | FIDYVLDKMD ELVQNTQRAL RVVEYLTETF PEKSVIGEKG EEVREELPEL<br>KEEFEKILER YKEIKENGPE SMEELYELYK ELREMYRKVY EAAKKVIESF<br>DWEKYPEVVE KVKELLAEEV PEK |
| dnAS9 | FIDYVYDKMD ELVQNTKRTL RVVEYLTETF PEKEVIGEEG AEVREKLPEV<br>KKEFEKILER YKKIKENGPK SLEELYELYK ELVEIYRKVY EMAKKVIRSF<br>DWEKHPEVVE EVRKLLAEEV PPE |
| dnAS10 | MSTRLPPRRR GEVMGVFNRV SREQGIDMGM HWRRMYEKVE QGTMTKATRG<br>EIMREYMVDM VDVAWRVKRH HPDIFEDMRE VMMRRTVRRY YRHPDNPELR<br>RIFRTMHMFL RTMFEKLAAI KIEPT |
| dnAS11 | MSRGLSEEEK EKVKEVIEKV AKELGIDLEK YFEELYKLIE EGKMTDESFK<br>KIMTEYFKDL VEVLWKIKKY YPEYFEKAEE VLLGDTVRLY YSHPDNPELQ<br>RFYKTLLYNL ELIFKKIRSI KIEEE |
| dnAS12 | MSRGLSEEEV EEVRKVFEEV AKELGIDLSK YFKELYDLIN EGKMTKESFK<br>EIMTKFNRDL VEVLWKIKVE HPEYFAKAER VLLGDIVKKY YSHPDNPEMQ<br>RFYKSLGYQL ELIFKRLRAI EIPEP |
| dnAS13 | MSLSAREQRW FDWYVHQMVV HGYEHWVGKV PEYAQNPEVS PARIEWMTKM<br>VEIWGRMINT MTPTQVSEVF IQWVIRHTHK FRFGDRVVEY VPTPTGTEDA<br>DGKMVYDPAS MQVERVRFQQ IMMREMHAYY NRS |
| dnAS14 | MLSEEEQKYF TYFIYLLVKH RLEKLKKELP VMEKDPSVPQ EKIDYKKKMV<br>ELWGKLLETK SDEEVANIFV EYIIRHESKF KFGDKVVTYK PKLTGTYDEN<br>GKFIYDKESV EKEKEEFKEL FLKEYKEYYE KK |
| dnAS15 | MLSEEEEEYF YYFVYLLVKH RLEKLKKELP VMKKDPSVSK EKIDYKEKLI<br>KLYGELLETK SNREVANTFI EWVIRHPVKF RFGDKVVTFE PKLTGTYDEN<br>GKFVYDKESV EKEKEKFKEL LLKEYKEYYE KK |
| dnAS16 | MSMTQKEVWK RFKRTMYEVD MDVYIFKRVY NMSRPYREEY YERMTAKYGI<br>TMPQEDIFDE IDRWPPVFDT MGTEMQRMKD IHKTTPWTRA WFVEFAGVMI<br>THMQGMMNMS PEVQRFVFGD PNPTERMRED VEIFTMVHDM WAG |
| dnAS17 | MHMSLEELWA KFRKALYEID MYVYIAKLVL DLKEEYFKEY YEIAKKKYGI<br>TTPYEEIKKE LEKLEPIINE IEEKLEELRE RHETEPFTRE WLVELLTGLI<br>ENVEKLLEIS EEVQKFIFGD ENPKERLKEI LEVLKAVRDH YGG |

*Continued on next page*

| Design | Sequence |
|---|---|
| dnAS18 | MSMSLEELWE KFRKALYRLD TYVYIFKKVY DLKEEYLEKY YEIRKKKYGI KMPFEKIKEE IEKLKPIIDE IGELLEELEE HHKTKPFTKE FLVELLSKLI EKVKELTKIS EEVQKFIFGD ENPKEELEEI IKVLEEVRNY YGG |
| dnAS20 | KEITPYDIFY FLDKFAKEVF GIDVREAMKD PEVRERMEKV WGYVYKNMEK AYKTKDEKEK EKYLFEAVEV FAQALLGRYL SEEEKEKIKK WLKENYEKYL IPK |
| dnAS21 | MDLSNAEIFH LFVYGMEKAY ENVETNPEEF ARYFLKSMYL LAKLREIVKT REFHGWLYDY LAPLVELAKK VYRGEEVPKE ELVKAWEPLY EKFKEIKPLI EKACTE |
| dnAS22 | MELSNAEIFH LFVYGMKKAA ENVETDPKKF AEYFLKSMYL LKKLREIVKT YEFHSFLYDK LAPLVELAKK VYKGEEVPKE ELLAAWEPLY KAFEEIKPII EKKCTE |
| dnAS23 | LSNKEIFNLF VYGMEKAHEN VESDPKKFAE YLLKSMYLLK KIREIVKTQE FHSFCYDYLA PLVELAKKVY KGEEVPKEEL LKAWEPLYEK FQEIKPIIEK KCTE |
| dnAS24 | MSELEEVLKA LDTVYEKLTA ALDTAQADPN ITLEQHRALV ETAWKAMVWM DEQVKKFTGL KKSLITIEHP KLFHEAYNAT SDLRAKLAQV RAAFEKTYAE AKKILEDYLK S |
| dnAS25 | MSELDKKIEA MEKIKEKLMK ALDTAIADPN ISKEEHQELV KTAWKFMVWL DDVVKEFTGE KKSLISIEDP KLFYEAYNAT DNFQEKLKEV KKRVEEVYEE AMKKLKEYKE S |
| dnAS26 | MSELERLYEF LKTFMSPEHQ EVFDKYYNKA KEAGSELLQY KVIGWFLLNY QSVRDGIEKY ISTLPEEKKA QFEKDWEVAL YFAHQIVDPG KEEYIPKFLA AMKPLVDYYF KEN |
| dnAS27 | MSKEEFLEAY KKVKEEWEKE FPKLVEELES GEISPEEAVE RLERFMRESA EYLWENSGPN MRHVRFTIEF LHAIATEMAR AIKRIRPDVA EEVRARIQEI YREIVRELLQ KYKDENYG |
| dnAS28 | MSREAFLEAY REVKKKWEEE FPKLEEKLRR GEISPEEAVE KLKEMMKESA KYLWENSGPN MRHVRFTIEF LHAMATRMAR AIKEIRPEVA EEVRERIQKE LREIVGELLE KYKKETEG |
| dnAS29 | MKVDLENFSE WSVKEKYEVA KKLHDIIREV GAKKEKEHPE YAKVVRLVTN WLCWFVDYVF KKSIENPSEE PLTPEKIFEE ARELFEKWIE RVAEKHGWSE EEKEFAREFF DEVLKAI |
| dnAS30 | KIDMEGYSEK SVKEKYKIAK EAHKVIREVG EEKIKEHPEY AKQFRLVENV ACWFFDYVFK NSIENPSSEP LTWEKIVKEF TELFEKWIER VAAKHGWSEE EKEFMREFFE EVLEKL |
| dnAS31 | EIDYSKEFPE LVEYFKELLY RAKKCMPPEY QKKFEEVIEI FEKGPFTYSM FLEMIEKLAE MFKDKEVGER FKEKAVALFK KLEELYKTDP EKFLEKFKRC FERLLWVLVQ LAKAEAK |
| dnAS32 | EIDLSKEFPE LVKYFLELLK KAQKCLPPKY QKKWEEVIKI FEEGPVTYEM FKEMIEKIAS MFEDKEVGEK FKEKALALYK KLEELYKTDP EKFLERFKRC FTRMLWVFVQ LARAESA |
| dnAS33 | MSFEEYVLDK MDELVQNTKR TARVIDYLTS TFPDKEVIGE EGAKVREEWP ALKKELEEIL KRYKEIKENG AESEEELYEL YKELREMYRK VYEAAKKVIN SFDWEKYPEV VEKVKELLKE EVPEE |
| dnAS34 | MSFEDYVLDK MDELVENTKR TLRVIEYLTE KFPDKEVIGE EGAKVREELP KVKEEFEKIL KEYEEIKENG AKSKEELYKL YEKLREMYRK VYEMAKKVIN SFDWEKYPEV IEEVKKLLSE EVPPE |
| dnAS35 | MSFLDYVFDK MDELVENTKR TLRVLEYLTS TFPDEKVIGK EGAKVKKELP ELKKEFEEIL KKYEKIKKNG PKSMEEAYKL YEELRELYRK VYEAAKKVIN SFDWEKYPEV IEKIKELLKE KVPEK |

*Continued on next page*

| Design | Sequence |
|---|---|
| dnAS36 | MELPEEIREK ARELTKLWLR IEYLGDKDLT PFNKLLEEVY NMEDVKEQFM LFARFVRSVL YGLKWLYEKF KGNEKEQEKV FEIIKKFPKF VEELYKKSKH KTKALYELVE KVKKECEEFI KKNKN |
| dnAS37 | MAMTLSPEEF AAQVLGDAYR VIKETKDVSK LEPYLDKIID YLSTYVDLPP KERALAVMYA VSYLTRLLVW LYQQGLISFE EAEEILLSFT KRLCEAMELD EKEQETVLNF TRRNIEKMKA WFEKHGR |
| dnAS38 | MAETLSPEEF SERVLGDAYR VIKETKDVSK LEPFLEKIIE YMSRYRDLPE RERALAAMHA VAYLTRLLVW LYQQGLISEE EMEELLRRFT ERLCEALELD EKSRATVMNF TDRNIAKARA WFAKHGR |
| dnAS39 | MDEEKKKILG KLSEGLLKLL RREYTKAYNA MEELDAKYRK ENGLPVDENS EPDPKIYNYC MDWFKERFEA QKKWLVKYFT ENYGSWGKVV EKLMDKLWEY FEEVMKTMPR REWTPENIID RLPIITEE |
| dnAS40 | MDEEKKEILR KLSIGLKNLL RKEWTKAYNA LKELDEKYRK ENNLPVDENS EYPEKIFNYC YDWFKERFEA QKKWLVKYFR ENYGSFGDVV EKLMERLWKE FEEVMKTMPP KDWTPENVID RLPIIDEE |
| dnAS41 | MGMSEVEKIE KLEELVEKLK ELAEKLKEVP EEERKKLREE MAKICREMLT LFQTLSVKSQ RYFLHKFWKE MKELIKENWS LLSEEEKEKT KALVKEIKAK YKGENVPLGY LTIMFTHLTG LDVTPEKSPT |
| dnAS42 | NKEVIIELFR CCYAIYEAVR TWSKDGYEGM KKLDEVYKNY LNNDEFYEKI ENSNIPEEIK TPFLALLNVT RYYMEEVLTN PEQRQGEAMK KLMRLANKFR YLAKQIYEKY PELKEEFEKI LKNPQVKL |
| dnAS43 | MSLSAWEVEV FKHFYELYNK HKETFESMPI EELYERLIKL FEEGIKRNEA LAKENPDDPR PRILAIIYRK LLEYIKKVKE KGEKDNKKFL EKLVAYLVEL LRKVKEVTGW SDEEIKEFAI NFLDEIFEGL PE |
| dnAS44 | LSEEERKRFT YHVYLLVKHR LEKLKKELPV MKKDPSVSKE KIEYKEKLIE LWGKLLETES DETVANTFVE YVIRNPFKFR FGDKVVTYNP KLTGTYDENG KFVYDKESVE KEKEEFEKLF MKEYKEYYEK K |
| dnAS45 | MFKEFMTKVY RATQRLFYVL KQFFEENNIV SKEEFEKFYK RVLYPILKMY YEEKVTPEDM EIVIEGYREL VELISKHVSK EDAEKLEEYF EAFEKAVRDI AENGIPEDPE EAQKYLNTWY GFCKTVYEVI TGK |
| dnAS46 | FEEFIEKLYR ATQRLFYVVE QYCEKNNIMS KEEFEKFKQK ILYPILKMYY EKKVTKEDIE IVIEGFEELV DLISKHVSEE DAEKLREYFE AFKEKLKEIA ENGIPEDPEK AQEYINYFYG FCGTVYRVIT GK |
| dnAS47 | KVRDEILEIY LEVYNEFVES FKPLFEGTGV ERLPEFELKT LEEIREFVNS LENPEDPMEV VKALHKRLVK TMKAALYMYL QLNPELKYPK DDEEKQKEFW EELKKTEFYK VYNSQVPKIL EAIKKMYYLA EKWRKENG |
| dnAS48 | DEARRKILEI YLEVHNEFVE SFKPLFEGTG VEILPEFQLK SLEEIEEEVE KLENPENPME VVKFLHERLV KTMKAALYMY LQMHPELKFP KDDEEAQKKW WEEFKKTELA KVYNSQIPKI LEAIKKMYYI AKQYEEENG |
| dnAS49 | GAAALDASRF APGESRREQM LRLLDDAIIE TLIEILELVT GKKYSPEEAL EFAKNSKDKT LKGLYLWYEE FRRLGEEFVD ALEARDYEKA REIVKKMYEN FLKMMNSVAV NKNLTPEQYR ELMRRGTARF REAVQAAIDL AL |
| dnAS50 | MPLSDAERFV RYLTEYWEEI EAMIEKQFAK LAEWARENNM PPEHVAKIPE VGKRMIEIIR ELLKEPEAKD PFVFYRKLIE GLSKALEEIF PGYISAEEAK DYFESSFWPL REELEKEYGP IPWHQFILYH LIYSTIELIA SLP |
| dnAS51 | MPLSRDERFV RYMTEYWEEI EEMIRKQFER VKKWAEENNM DPEHVAKIPE VGEELIKIIK ELMEKPEAKD FRVFRRKLIE GLSKMLEKTF PGFISAEEVY NYFVSSYWPL REELEKEYGP IPEFRFFLYH FIYTTIEIIA SLP |

*Continued on next page*

*Supplementary Table S1 (continued)*

| Design | Sequence |
|---|---|
| dnAS52 | TVTITTPDGK EVTVEVAYGR RDLGDGVVRY YFKVNDVTIE IDINKVTKKV TLSLYLKPPT DGESYVDKIF TAVWYIIKYF ESKGVNVEGA YKELKELLEK FKNGEIEETP YAIWKACAEL LLNFFRRNKI LPEEMLENTE FKPPKVLV |
| dnAS53 | MKSNYEIFEE IYNKYKPKIE KIYELYEKGY ISKEEYYELT NKVLKYMFYE IYKKTGIRYD YYLYEELQKK KEKKLSFKEA LELALEEILK KLKEEDK |

Supplementary Table S2: **Expression and binding properties of designed $\alpha$-synuclein binders.** Expression status, reported screening outcomes and SPR affinities are listed for all 52 designs. Unresolved expression entries have incomplete or conflicting records. N.D., affinity not determined; N.R., no assay record available. An undetermined affinity does not imply absence of binding. Reported $K_D$ values are retained pending final source-fit verification.

| Design | Expressed | Assay | Binding result | $K_D$ (nM) |
|---|---|---|---|---|
| dnAS1 | No | N.R. | Not detected | N.D. |
| dnAS2 | No | N.R. | Not detected | N.D. |
| dnAS3 | Yes | SPR | Not detected | N.D. |
| dnAS4 | No | N.R. | Not detected | N.D. |
| dnAS5 | No | N.R. | Not detected | N.D. |
| dnAS6 | Unresolved | N.R. | Not detected | N.D. |
| dnAS7 | No | N.R. | Not detected | N.D. |
| dnAS8 | No | N.R. | Not detected | N.D. |
| dnAS9 | Yes | SPR | Not detected | N.D. |
| dnAS10 | No | N.R. | Not detected | N.D. |
| dnAS11 | No | N.R. | Not detected | N.D. |
| dnAS12 | Yes | SPR | Not detected | N.D. |
| dnAS13 | No | N.R. | Not detected | N.D. |
| dnAS14 | No | N.R. | Not detected | N.D. |
| dnAS15 | Yes | SPR | Binding | 211 |
| dnAS16 | No | N.R. | Not detected | N.D. |
| dnAS17 | Yes | N.R. | Not detected | N.D. |
| dnAS18 | Yes | SPR | Not detected | N.D. |
| dnAS20 | Yes | SPR | Not detected | N.D. |
| dnAS21 | Yes | SPR | Not detected | N.D. |
| dnAS22 | Yes | SPR | Binding | 13000 |
| dnAS23 | Yes | SPR | Not detected | N.D. |
| dnAS24 | Yes | SPR | Not detected | N.D. |
| dnAS25 | Unresolved | N.R. | Not detected | N.D. |
| dnAS26 | Yes | SPR | Not detected | N.D. |
| dnAS27 | Yes | SPR | Not detected | N.D. |
| dnAS28 | Yes | SPR | Not detected | N.D. |
| dnAS29 | Unresolved | SPR | Not detected | N.D. |
| dnAS30 | Unresolved | SPR | Not detected | N.D. |
| dnAS31 | Yes | SPR | Not detected | N.D. |
| dnAS32 | Yes | SPR | Not detected | N.D. |
| dnAS33 | Yes | SPR | Not detected | N.D. |
| dnAS34 | Yes | SPR | Not detected | N.D. |
| dnAS35 | Yes | SPR | Not detected | N.D. |
| dnAS36 | Yes | SPR | Binding | 566 |
| dnAS37 | Yes | SPR | Binding | 20000 |
| dnAS38 | Yes | SPR | Binding | 5300 |
| dnAS39 | Yes | SPR | Not detected | N.D. |
| dnAS40 | Yes | SPR | Not detected | N.D. |
| dnAS41 | Unresolved | N.R. | Not detected | N.D. |
| dnAS42 | Yes | SPR | Binding | 1300 |
| dnAS43 | Unresolved | SPR | Not detected | N.D. |
| dnAS44 | Unresolved | SPR | Not detected | N.D. |

*Continued on next page*

*Supplementary Table S2 (continued)*

| Design | Expressed | Assay | Binding result | $K_D$ (nM) |
|---|---|---|---|---|
| dnAS45 | Unresolved | SPR | Not detected | N.D. |
| dnAS46 | Yes | SPR | Not detected | N.D. |
| dnAS47 | Yes | SPR | Not detected | N.D. |
| dnAS48 | Yes | SPR | Not detected | N.D. |
| dnAS49 | Unresolved | SPR | Not detected | N.D. |
| dnAS50 | Unresolved | SPR | Binding | 10000 |
| dnAS51 | Yes | SPR | Not detected | N.D. |
| dnAS52 | Yes | SPR | Not detected | N.D. |
| dnAS53 | Yes | SPR | Not detected | N.D. |

Supplementary Table S3: **Amino-acid sequences of designed RBX1 binders.** Sequences of the 12 designs are listed in blocks of ten residues. Design identifiers correspond to Supplementary Table S4; original competition names are provided where applicable.

| Design | Original name | Sequence |
|---|---|---|
| dnRB1 | deep-bear-clay | DRAALEALGK EVLERVKGLV DDLVKVMNDP DASRMEIAEA ARKMVSETAR IAGEYAAKIP<br>AGSAPTAVDY RLYEEMMLGF LIGVTVRVIL EKEGYGLAPT AEEVRGFVTN FTRWMKRFPA<br>LPRKYLAKAA VIVRWTVRQV PLEGSGLTEE EIRELSEGFP ASFGIDIETD PEVKVIIHEL<br>VTNVPEELKE AQKEFPLIKD FYDYLRELLK KIES |
| dnRB2 | jade-cobra-cypress | SAEKEAELEK EKEKILEKVK EVNPDFSEEQ SETIAESIFT MNYYIASGNP ESEKVINAGA<br>KIRKILYFDK PVNEEQLKKI KEVYLESFKK IYEILHPELK L |
| dnRB3 | jade-mole-cypress | PYISEEARKK VESTIATLRE QKEALRKAGY PEAAELTEKA IATLEELAEK MRKEREIPTE<br>EKLERYCEFL ELMVEYYRYA LEAGEEVQKA AVEISIQMYA GDNPVRQAAR KEYEDPANSL<br>REKVERAYYE TKVGLKIISE LLEYYKKKYE ESGGKYYPTL EELEKQFEEA NKKYEEKREE<br>LYEEYREEVG |
| dnRB4 | noble-gecko-sand | MTRMELYEAG RNLTPEEVKS RLAESREIIR EGHAIVTNLT PEEFTSWEEY YKFHSIYRQV<br>QQISIMVLAH VTENMVVTEE NRDMYRMMRF IADVEIDPWA NWPQVQKLME KASPKYMTIP<br>WSPQHWWMRH VFHMVSMGAM IRVVEEMEVT DGKWVWTAAS MEAFEVFLEI MEMRMEAYDK<br>FLTGKDWYTT MRGMIAKEQA M |
| dnRB5 | noble-gecko-bronze | STYEDLNGLI DIYEENPADP RKAEEKIGEY LDEAIKRNPD NKVIEKLKET YETHKRYYTT<br>RENPEGYYTE YYSTQRELAK ELAKEVDKEK LEEVVERTYE TRDKMAEACE KAKENPEDYE<br>RLASYEAETT YTYIEAMSEQ DPSVISRYRE KLAARAEEHK EKAEKEKSAH GYETVVTLRT<br>QVVQLEISEI YDEAEKAGKS PEDAEKEAAK AQAAAKPAIE AERQKSAQLY YETL |
| dnRB6 | radiant-falcon-rose | MITVDEIYEI HKSGNKKEAF SAHADTLFES MRARMANGAA AGNLDFEDMF VHMYVTSMYY<br>YYPEHVTAAQ RTIVAETVGT MVNVLENSAY PLVGQLIRQV TATAPDAMSW YERHDKRHII<br>AFAFLYYLWR WLEANDMYLT PKMEEWLKWV AKTIFHQLMM WELDEDVTRI LTEMASEMPF IEVM |
| dnRB7 | vast-goat-ember | MWHMTPMQRQ VRVVLYQPEN MALLMRFMDE WLEKHVANPN NPFRPQFAAM VPHWGVHKTV<br>YNKHADFIAY RYKNPNGPPP SMSGAERTEK EGAWETFQEH WHQFKHDAYK VLTKMFPPLR<br>ALVDIHSDMM MEMWGKTPGE RDDIEEAYYD QIVTEMKQFI RQLKPHVSPA VASEMKIEV |
| dnRB8 | – | KAKKLYEMLR KGRDPEKRDE FPKLFYEVFG EIDQSEFLPF FEKLKPYFRK MVEALASRID<br>PAKYPGLGEV LEELRRILDD PNPSVVDFFS ALLRLWGELA RYKAFISEDN PEGDFEVIFE<br>SLLYLNKGYE EAFKSLKEPL SSSLLELLEV LLLLNAFGFF AAAYYYHKKT GEPPETLVPK<br>LREVWEASLR ALETAYRKGN FSEEEIELKR EIFEELLEEN IELFILLHDP ENRHKAFEKL<br>YKWKKEIEE |
| dnRB9 | – | MPPISVENLL HFIIGHVMEF VDPSEEEKVL ERLAEKFGKT LEEFMKQFYE ELAQQPEEFR<br>ELLKLLEEAK NSIDAEAVFK KDPALVKRRE EVVKRVFELL DTLSEPLSDE ELIDKFTLET<br>LKLLLETLKT VKNSNLSEEE KKELLMLISM EACLFIIARH VAGIREIPSE IQKIFLEILE<br>LAKEEIA |
| dnRB10 | – | MKIQPVTDED VEAVWEEIYP IFELLHKLSK MSEEEKKSWL KENWDDFLNK VKAVFDKLFE<br>ICRRSDDYEG ILKLLYKLKD FLKNGPPGGP EALPFSEEKR QKLVDFIDLI IEIIESNPEL<br>SVRNKKCLVI LAFLVIFTGI LNEVKKYVIE TEEEWLSFHA SLHKVAQRIV DFVLNLRKGE<br>EISLDEVEEL WNLLFEGFK |
| dnRB11 | – | PEFSEEVKPL LEKFLKDFLK YQPYLDLLYE ALRRAINLLA FYGINDPKGV EEAKKYLEDV<br>VKNGTPEQQA WARVAIVIIE LLAKRKDPII TEALEKHSKL LKTPYTGTKE ELEEYFEVLK<br>ELLEAEIFFL ESLSKEELEK LGKAIGESPH PGIKRVLSPE EMVHVLSVLL LVYRLALELL<br>DWYVENGYVL DWEAVKEVAY KNLEKYLEKV KEWLEKKLKE LSE |
| dnRB12 | – | MSPEVREKLA SLPKEVMEHN VEVKIVLHVA FCLLIHQMRK ALEAGTEITF PKEVELMLRA<br>ASMMFFPGNY EAQLALRDLT DRLLEALRAR DYAEAIRLLA EFEKTLLTYL TGSGRPLSEE<br>ELRSFYGGRT PEEIAEILVE AVERGELTPE QALAIWVVID AHHWRELYEE LGYLTEEQRE<br>WFETVIEFCE VVFNIDVEEA IGIR |

Supplementary Table S4: **Expression and binding properties of designed RBX1 binders.** Expression outcomes and binding results are listed for 12 designs. Affinities were obtained from 1:1 kinetic fits of SPR or BLI measurements.

| Design | Expressed | Assay | Binding result | $K_D$ (nM) |
|---|---|---|---|---|
| dnRB1 | Yes | SPR | Binding | 85 |
| dnRB2 | Yes | SPR | Binding | 170 |
| dnRB3 | Yes | SPR | Not detected | N.D. |
| dnRB4 | Yes | SPR | Not detected | N.D. |
| dnRB5 | Yes | SPR | Not detected | N.D. |
| dnRB6 | Yes | SPR | Not detected | N.D. |
| dnRB7 | No | – | Not expressed | N.D. |
| dnRB8 | Yes | BLI | Binding | 4.5 |
| dnRB9 | Yes | BLI | Binding | 240 |
| dnRB10 | Yes | BLI | Binding | 169 |
| dnRB11 | Yes | BLI | Not detected | N.D. |
| dnRB12 | Yes | BLI | Not detected | N.D. |

N.D., affinity not determined. dnRB7 was not expressed and is not classified as a binding negative.

Supplementary Table S5: **Amino-acid sequences of designed intracellular nanobodies against $\beta_1$AR.** VHH sequences of the 24 designs are listed alongside Nb6B9 and Nb80 reference sequences in blocks of ten residues. Vector-encoded affinity tags and cleavage sites are excluded. Design identifiers correspond to Supplementary Table S6.

| Design | Sequence |
|---|---|
| Nb6B9 | QVQLQESGGG LVQAGGSLRL SCAASGSIFA LNIMGWYRQA PGKQRELVAA IHSGGTTNYA NSVKGRFTIS RDNAANTVYL QMNSLKPEDT AVYYCNVKDF GAIIYDYDYW GQGTQVTVSS |
| Nb80 | QVQLQESGGG LVQAGGSLRL SCAASGSIFS INTMGWYRQA PGKQRELVAA IHSGGSTNYA NSVKGRFTIS RDNAANTVYL QMNSLKPEDT AVYYCNVKDY GAVLYEYDYW GQGTQVTVSS |
| NbIC1 | QVQLQESGGG LVQAGGSLRL SAAASGGIYV LNVMGWYRQA PGKQRELVAA THTKGMTNYA NSVKGRFTIS RDNAANTVYL QMNSLKPEDT AVYYGYVKDY GNLIWDEIYW GQGTQVTVSS |
| NbIC2 | QVQLQESGGG LVQAGGSLRL SAAASGGIHV LNVMGWYRQA PGKQRELVAA THSNGETNYA NSVKGRFTIS RDNAANTVYL QMNSLKPEDT AVYYGYVKDY GNITYTETYW GQGTQVTVSS |
| NbIC3 | QVQLQESGGG LVQAGGSLRL SAAASGGIHV LNVMGWYRQA PGKQRELVAA THSNGSTNYA NSVKGRFTIS RDNAANTVYL QMNSLKPEDT AVYYGYVKDY GNITYVETYW GQGTQVTVSS |
| NbIC4 | QVQLQESGGG LVQAGGSLRL SAAASGGIFA LNVMGWYRQA PGKQRELVAA ADYNNVTNYA NSVKGRFTIS RDNAANTVYL QMNSLKPEDT AVYYGYVQDY GQVIYDDIYW GQGTQVTVSS |
| NbIC5 | QVQLQESGGG LVQAGGSLRL SAAASGGIHP LNVMGWYRQA PGKQRELVAA ADYKDNTNYA NSVKGRFTIS RDNAANTVYL QMNSLKPEDT AVYYGYVKDF GNITYTDTYW GQGTQVTVSS |
| NbIC6 | QVQLQESGGG LVQAGGSLRL SAAASGGIHP LNVMGWYRQA PGKQRELVAA ADYNDNTNYA NSVKGRFTIS RDNAANTVYL QMNSLKPEDT AVYYGYVKDF GNITYTDTYW GQGTQVTVSS |
| NbIC7 | QVQLQESGGG LVQAGGSLRL SAAASGSIFV LQVMGWYRQA PGKQRELVAA IHSKGTTNYA NSVKGRFTIS RDNAANTVYL QMNSLKPEDT AVYYAYVKDF GAIVYDEYYW GQGTQVTVSS |
| NbIC8 | QVQLQESGGG LVQAGGSLRL SAAASGAIHV LNVMGWYRQA PGKQRELVAA IHSNGETNYA NSVKGRFTIS RDNAANTVYL QMNSLKPEDT AVYYAYVKDF GNITYVEEHW GQGTQVTVSS |
| NbIC9 | QVQLQESGGG LVQAGGSLRL SAAASGAIHV LNVMGWYRQA PGKQRELVAA IHSNGETNYA NSVKGRFTIS RDNAANTVYL QMNSLKPEDT AVYYAYVEDF GNITYTEKYW GQGTQVTVSS |
| NbIC10 | QVQLQESGGG LVQAGGSLRL SAAASGGIYV LNVMGWYRQA PGKQRELVAA IHSKGDTNYA NSVKGRFTIS RDNAANTVYL QMNSLKPEDT AVYYSYVQDF GNVIYDEIYW GQGTQVTVSS |
| NbIC11 | QVQLQESGGG LVQAGGSLRL SAAASGGVHV LNVMGWYRQA PGKQRELVAA IHSNGETNYA NSVKGRFTIS RDNAANTVYL QMNSLKPEDT AVYYSYVKDY GNITYTVTTW GQGTQVTVSS |
| NbIC12 | QVQLQESGGG LVQAGGSLRL SAAASGGVHV LNVMGWYRQA PGKQRELVAA IHSNGETNYA NSVKGRFTIS RDNAANTVYL QMNSLKPEDT AVYYSYVKDY GNITYVETYW GQGTQVTVSS |

*Continued on next page*

*Supplementary Table S5 (continued)*

| Design | Sequence |
|---|---|
| NbIC13 | QVQLQESGGG LVQAGGSLRL SAAASGAIYV LNVMGWYRQA PGKQRELVAA EHSLGETNYA NSVKGRFTIS RDNAANTVYL QMNSLKPEDT AVYYAYVKDF GAMTYDEVYW GQGTQVTVSS |
| NbIC14 | QVQLQESGGG LVQAGGSLRL SAAASGGIHV LNVMGWYRQA PGKQRELVAA EHSLGETNYA NSVKGRFTIS RDNAANTVYL QMNSLKPEDT AVYYAYVKDF GEITYTDTYW GQGTQVTVSS |
| NbIC15 | QVQLQESGGG LVQAGGSLRL SAAASGGIHV LNVMGWYRQA PGKQRELVAA EHSLGETNYA NSVKGRFTIS RDNAANTVYL QMNSLKPEDT AVYYAYVKDY GEITYVSTSW GQGTQVTVSS |
| NbIC16 | QVQLQESGGG LVQAGGSLRL SAAASGGIYV LNVMGWYRQA PGKQRELVAA IASNGVTNYA NSVKGRFTIS RDNAANTVYL QMNSLKPEDT AVYYSYVKDY GNLVYDEIYW GQGTQVTVSS |
| NbIC17 | QVQLQESGGG LVQAGGSLRL SAAASGGVHV LNVMGWYRQA PGKQRELVAA IASDGTTNYA NSVKGRFTIS RDNAANTVYL QMNSLKPEDT AVYYSYVKDY GNITYDVTYW GQGTQVTVSS |
| NbIC18 | QVQLQESGGG LVQAGGSLRL SAAASGGIHV LNVMGWYRQA PGKQRELVAA IASDGTTNYA NSVKGRFTIS RDNAANTVYL QMNSLKPEDT AVYYSYVKDY GEITYDVTYW GQGTQVTVSS |
| NbIC19 | QVQLQESGGG LVQAGGSLRL SAAASGGVFV LNVMGWYRQA PGKQRELVAA IHSSGDTNYA NSVKGRFTIS RDNAANTVYL QMNSLKPEDT AVYYSYVIDF TNIIWDDVYW GQGTQVTVSS |
| NbIC20 | QVQLQESGGG LVQAGGSLRL SAAASGGVHV LNVMGWYRQA PGKQRELVAA IHSSGETNYA NSVKGRFTIS RDNAANTVYL QMNSLKPEDT AVYYSYVDDF TNITYTETYW GQGTQVTVSS |
| NbIC21 | QVQLQESGGG LVQAGGSLRL SAAASGGVHV LNVMGWYRQA PGKQRELVAA IHSSGETNYA NSVKGRFTIS RDNAANTVYL QMNSLKPEDT AVYYSYVDDY TNITYTNTYW GQGTQVTVSS |
| NbIC22 | QVQLQESGGG LVQAGGSLRL SAAASGGIFI LNTMGWYRQA PGKQRELVAA IHSLGDTNYA NSVKGRFTIS RDNAANTVYL QMNSLKPEDT AVYYAQVVDY GQVTYDEIYW GQGTQVTVSS |
| NbIC23 | QVQLQESGGG LVQAGGSLRL SAAASGGVHV LNTMGWYRQA PGKQRELVAA IHSNGETNYA NSVKGRFTIS RDNAANTVYL QMNSLKPEDT AVYYAQLDDY GEITYVNRSW GQGTQVTVSS |
| NbIC24 | QVQLQESGGG LVQAGGSLRL SAAASGGVHV LNTMGWYRQA PGKQRELVAA IHSNGETNYA NSVKGRFTIS RDNAANTVYL QMNSLKPEDT AVYYAQVDDY GNITYTERSW GQGTQVTVSS |

Supplementary Table S6: **Expression and binding properties of designed intracellular nanobodies against $\beta_1$AR.** Soluble expression yields, interface hydrophobicity and apparent affinities are listed for 16 GuideFlip designs and eight hallucination-only controls. Binding to isoprenaline-bound $\beta_1$AR was measured by BLI.

| Design | Expression yield (mg/50 mL) | Interface hydrophobicity (%) | Assay | Binding result | $K_D$ (nM) |
|---|---|---|---|---|---|
| NbIC1[a] | 0.009 | 44.8 | BLI | Not detected | N.D. |
| NbIC2 | 0.024 | 39.1 | BLI | Not detected | N.D. |
| NbIC3 | 0.026 | 36.4 | BLI | Not detected | N.D. |
| NbIC4[a] | 0.008 | 55.2 | BLI | Not detected | N.D. |
| NbIC5 | 0.042 | 44.0 | BLI | Binding | 45 |
| NbIC6 | 0.088 | 45.5 | BLI | Binding | 13 |
| NbIC7[a] | 0.015 | 45.2 | BLI | Binding | N.D. |
| NbIC8 | 0.095 | 41.7 | BLI | Binding | N.D. |
| NbIC9 | 0.021 | 41.7 | BLI | Not detected | N.D. |
| NbIC10[a] | 0.035 | 44.8 | BLI | Binding | 8 |
| NbIC11 | 0.065 | 33.3 | BLI | Binding | N.D. |
| NbIC12 | 0.157 | 38.1 | BLI | Binding | N.D. |
| NbIC13[a] | 0.019 | 50.0 | BLI | Binding | 200 |
| NbIC14 | N.D.[b] | N.R. | N.R. | Not recorded | N.D. |
| NbIC15 | 0.022 | 43.5 | BLI | Binding | N.D. |
| NbIC16[a] | 0.070 | 48.3 | BLI | Binding | 440 |
| NbIC17 | 0.396 | 45.8 | BLI | Binding | N.D. |
| NbIC18 | 0.092 | 45.5 | BLI | Binding | N.D. |
| NbIC19[a] | 0.023 | 53.6 | BLI | Binding | $<1$ |
| NbIC20 | 0.091 | 38.5 | BLI | Binding | 21 |
| NbIC21 | 0.289 | 38.5 | BLI | Binding | N.D. |
| NbIC22[a] | 0.039 | 44.8 | BLI | Binding | 96 |
| NbIC23 | 0.133 | 40.9 | BLI | Binding | N.D. |
| NbIC24 | 0.055 | 34.8 | BLI | Binding | N.D. |

[a] Hallucination-only control; all other designs used GuideFlip. [b] NbIC14 was not expressed. N.D., affinity not determined; N.R./not recorded, no record available. For binders with N.D. affinity, no reliable global fit was obtained.

Supplementary Table S7: Weights of the AlphaFold design-loss terms used by GuideFlip.

| Term | Weight |
|---|---|
| Binder confidence (pLDDT) | 0.1 |
| Interface confidence (iPTM) | 0.05 |
| PAE within binder | 0.4 |
| Interface PAE (iPAE) | 0.1 |
| Contacts within binder | 1.0 |
| Contacts binder–target | 1.0 |
| Radius of gyration | 0.3 |
| Helicity | $-0.3$ (helix) / $-2$ ($\beta$) / 0.0 (nanobody) |

**Algorithm 1** GuideFlip sampling

**def** GuideFlip($\texttt{target}, L; T, w, \eta_g, \eta, \beta, \delta, \tau_0, \tau_1$):

$\ell_t \leftarrow$ random logits; $\quad s \leftarrow \text{MASK}^L$; $\quad \mathscr{C} \leftarrow \emptyset$
$p_t \leftarrow \text{softmax}(\ell_t)$; $\quad t \leftarrow 0$; $\quad \lambda_t \leftarrow 0$
**repeat**
  **if** $\lambda_t \neq 0$ **then**
    $\ell_{\text{ADFlip}} \leftarrow \text{ADFlip}(X_t, s_{\mathscr{C}})$
    $u \leftarrow \log p_t + \ell_{\text{ADFlip}}/T$
    $X', \nabla_u \mathscr{L}_{\text{AF}} \leftarrow \text{AF}(\text{softmax}(u))$
    $g \leftarrow -\eta_g \dfrac{\nabla_u \mathscr{L}_{\text{AF}}}{\|\nabla_u \mathscr{L}_{\text{AF}}\|_{\text{rms}}}$
    $p_{t+\Delta t} \leftarrow \text{softmax}(u+g)$
    $X_{t+\Delta t} \leftarrow X'$ **if** $\text{iRMSD}(X', X_{\text{best}}) \leq \delta$ **else** $X_t$
    $(s, \mathscr{C}) \leftarrow \text{decode}(p_{t+\Delta t}, s, \mathscr{C}, \tau(t))$
    $(p_t, X_t) \leftarrow (p_{t+\Delta t}, X_{t+\Delta t})$
  **else**
    $u \leftarrow \beta p_t + (1-\beta)\ell_t$
    $X_t, \nabla_{\ell_t} \mathscr{L}_{\text{AF}} \leftarrow \text{AF}(u)$
    **if** first evaluation or confidence reaches a new best **then**
      $(p_{\text{best}}, X_{\text{best}}) \leftarrow (p_t, X_t)$
    **end if**
    $\ell_t \leftarrow \ell_t - \eta\sqrt{L}\dfrac{\nabla_{\ell_t} \mathscr{L}_{\text{AF}}}{\|\nabla_{\ell_t} \mathscr{L}_{\text{AF}}\|_2}$
    **if** confidence stops improving **then**
      $\ell_t \leftarrow (w/T)\log p_{\text{best}}$; $\quad X_t \leftarrow X_{\text{best}}$; $\quad \lambda_t \leftarrow 1$
    **end if**
    $p_t \leftarrow \text{softmax}(\ell_t)$
  **end if**
  $t \leftarrow \text{estimate\_time}(p_t)$
**until** $|\mathscr{C}| = L$
**return** $s, X_t$

**Notation.** $L$ is the binder length; $i$ indexes binder positions and $a \in \mathscr{A}$ the 20 amino acids. Sequence-score and probability arrays have shape $L \times 20$; softmax and argmax act over $a$. The input `target` contains the target sequence, structure and templates. Confidence denotes smoothed binder pLDDT.

| | |
|---|---|
| $s$, $\mathscr{C}$ | Partial sequence and assigned positions; $s_i = \text{MASK}$ for $i \notin \mathscr{C}$. |
| $\ell_t$, $\ell_{\text{ADFlip}}$ | Current binder logits and ADFlip logits. |
| $p_t$, $p_{t+\Delta t}$, $p_{\text{best}}$ | Current and updated probabilities, and the best saved distribution. |
| $u$, $g$ | Binder input scores and the current gradient-guidance step. |
| $X_t$, $X_{t+\Delta t}$ | Current complex and accepted complex for the next iteration. |
| $X'$, $X_{\text{best}}$ | Candidate complex and the best saved complex, fixed as reference after the switch. |
| $t$, $\Delta t$ | Time estimated from current probabilities and its change per update. |
| $\lambda_t$ | Prior gate (0: off; 1: on). |
| $T$, $w$ | Temperature and initialization-prior weight. |
| $\eta_g$, $\eta$, $\beta$, $\delta$ | Guidance scale, gradient step size, input blend and interface RMSD tolerance. |
| $\tau(t)$ | Decoding threshold: $\tau_0 + (\tau_1 - \tau_0)t$. |
| $\mathscr{L}_{\text{AF}}$ | AF design loss; target conditioning and backpropagation are implicit. |
| decode | Assign argmax residues at unassigned positions with $\max_a p_{ia} \geq \tau$; update $s, \mathscr{C}$. |

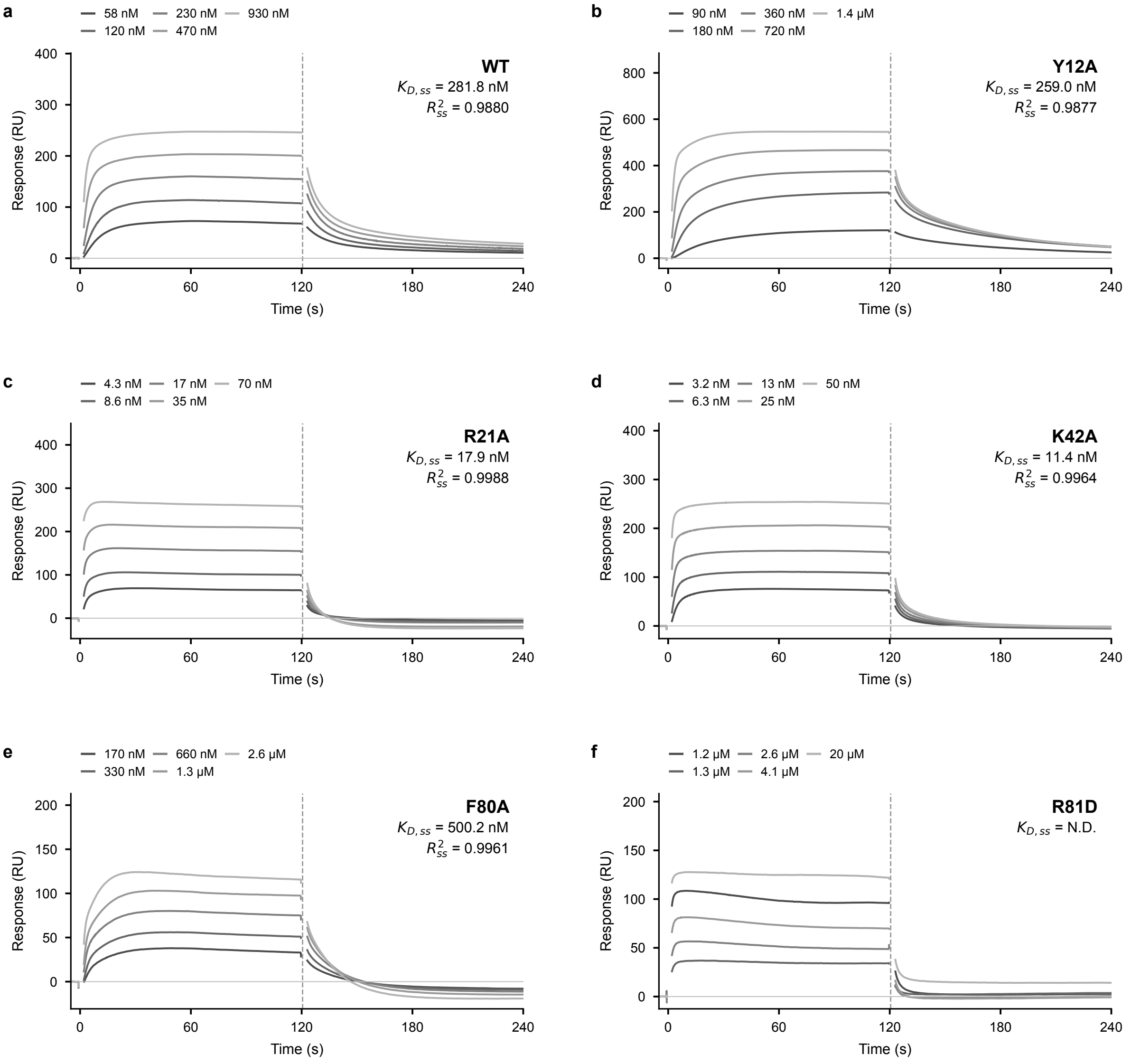


Supplementary Figure S1: **SPR sensorgrams of dnAS15 single mutants. a–f**, WT, Y12A, R21A, K42A, F80A and R81D. Grey curves show experimental responses; $K_{\mathrm{D,ss}}$ and $R^2_{\mathrm{ss}}$ are from 1:1 steady-state fits. One measurement per construct; WT and mutants were measured in separate runs. N.D., affinity not determined. R81D concentrations remain unverified.

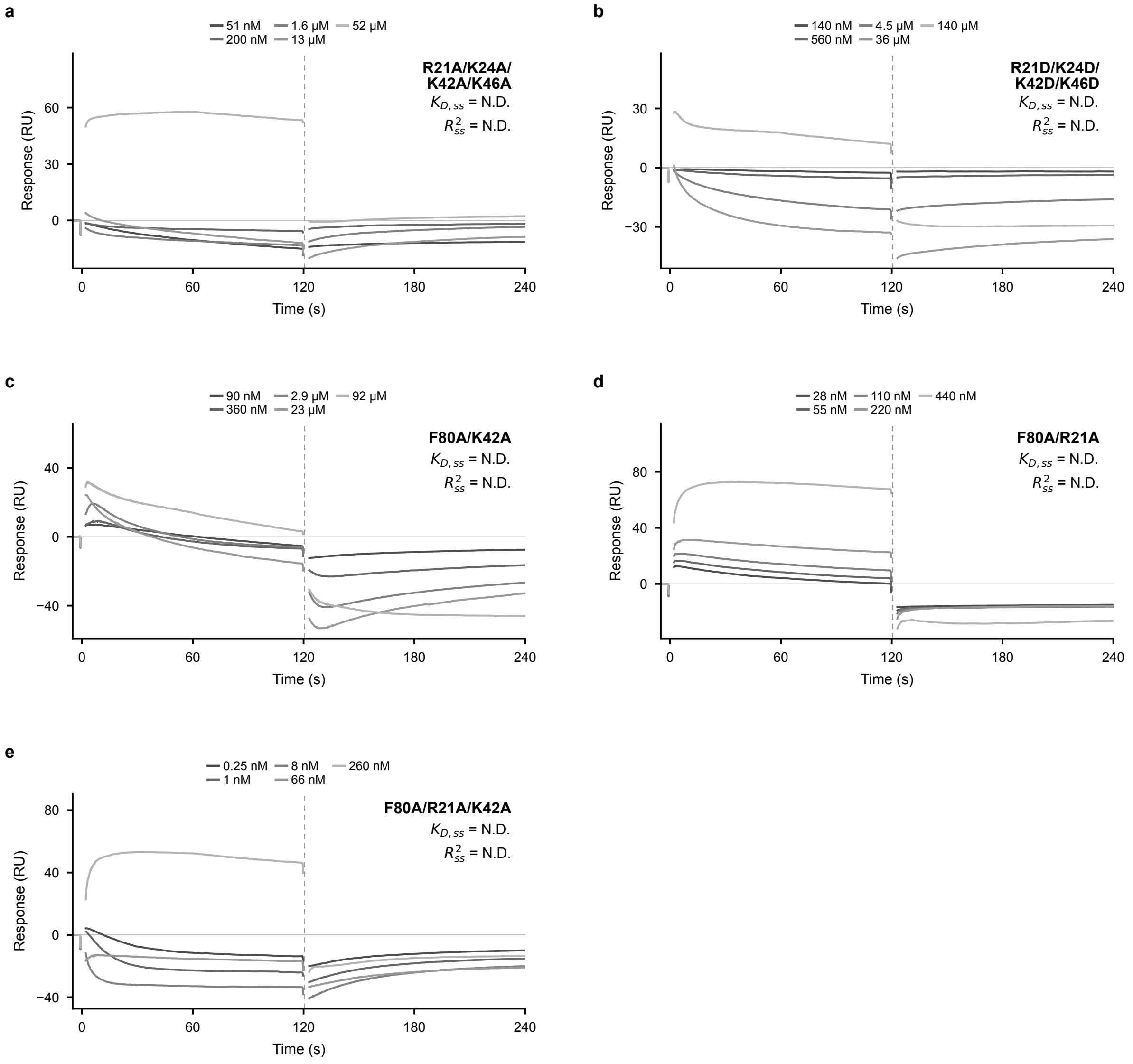


Supplementary Figure S2: **SPR responses of dnAS15 combination mutants. a–e**, R21A/K24A/K42A/K46A, R21D/K24D/K42D/K46D, F80A/K42A, F80A/R21A and F80A/R21A/K42A. Grey curves show experimental responses; no kinetic fits are shown. N.D., affinity not determined. Negative responses alone do not establish absence of binding.

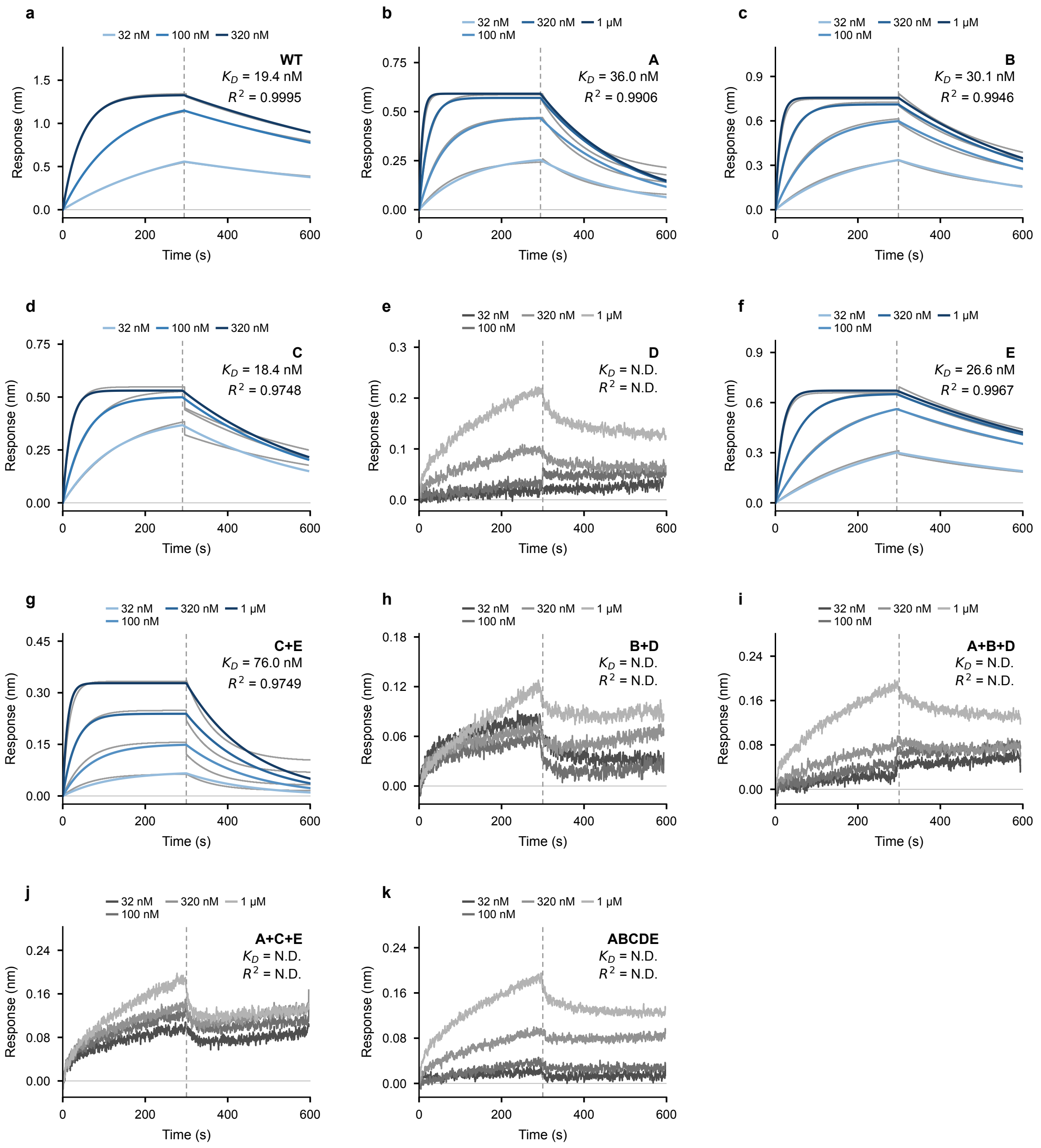


Supplementary Figure S3: **BLI binding measurements of dnRB1 interface mutants. a**, WT. **b–k**, A, B, C, D, E, C+E, B+D, A+B+D, A+C+E and ABCDE. Grey curves show experimental responses; blue curves show 1:1 kinetic fits. One measurement is shown per mutant at the indicated concentrations. N.D., affinity not determined. The recorded transition (approximately 300 s) differs from the 220-s association time in the protocol description.
